# RPG interacts with E3-ligase CERBERUS to mediate rhizobial infection in *Lotus japonicus*

**DOI:** 10.1101/2022.06.30.498293

**Authors:** Xiaolin Li, Miaoxia Liu, Min Cai, David Chiasson, Martin Groth, Anne B. Heckmann, Trevor L. Wang, Martin Parniske, J. Allan Downie, Fang Xie

## Abstract

Symbiotic interactions between rhizobia and legumes result in the formation of root nodules, which fix nitrogen that can be used for plant growth. Rhizobia usually invade legume roots through a plant-made tunnel-like structure called an infection thread (IT). Rhizobium-directed polar growth (*RPG*) encodes a coiled-coil protein that was identified in *Medicago truncatula* as required for root nodule infection, but the function of RPG remains poorly understood. In this study, we identified and characterized *RPG* in *Lotus japonicus* and determined that it is required for IT formation. *RPG* was induced by *Mesorhizobium loti* or purified Nodulation factor and displayed an infection-specific expression pattern. Nodule inception (NIN) bound to the *RPG* promoter and induced its expression. A GFP-RPG protein was localized in puncta subcellular localization in *L. japonicus* root protoplasts and in root hairs infected by *M. loti*. The N-terminal predicted C2 lipid-binding domain of RPG was not required for this subcellular localization or for function. CERBERUS, a U-box E3 ligase which is also required for rhizobial infection, was found to be localized in similar puncta. RPG co-localized and directly interacted with CERBERUS at the early endosomes (TGN/EE) compartment and near the nuclei in root hairs after rhizobia inoculation. Our study sheds light on that a RPG-CERBERUS protein complex that is involved in an exocytotic pathway mediating IT polarity growth which is driven by nuclear migration.

**One sentence summary:** Puncta localization RPG-CERBERUS protein complex promote polarity growth of ITs driven by nuclear migration.

## INTRODUCTION

Nitrogen-fixing root nodule symbioses (RNS) between legumes and their rhizobial symbionts are important because the plants can obtain nitrogen from gaseous N_2_ reduced to NH_3_ by rhizobia. The establishment and maintenance of this symbiosis depend on a molecular dialogue between the partners. The formation of N_2_-fixing nodules requires two developmental processes: nodule organogenesis and bacterial infection. Although the two processes can be genetically separated, they must be spatially and temporally coordinated to ensure nodule organogenesis at sites of bacterial infection (Oldroyd and Downie, 2008). In response to flavonoids exuded by the plant, rhizobia secrete decorated lipochito-oligosaccharide molecules called nodulation factors (NFs) that can activate nodule organogenesis, and can induce cellular changes associated with the initiation of bacterial infection (Oldroyd and Downie, 2004).

In most legume-rhizobium symbioses, rhizobia invade legume roots *via* root hairs. Rhizobia attach to root hairs, triggering root-hair curling that entraps rhizobia which induce localized cell-wall degradation and rearrangement of the plant cytoskeleton, contributing to the formation of plant-made tunnel-like structures called the infection threads (ITs) (Gage, 2004). Rhizobia colonize the ITs which grow through root cells, ultimately reaching the nodule primordium; the bacteria are then budded off surrounded by a plant-made membrane into plant cells in which they fix nitrogen using carbon supplied by the plant (Robertson et al., 1978). Genetic studies in *Lotus japonicus* and *Medicago truncatula* have identified several genes required for IT initiation and formation. Some are associated with changes in the actin cytoskeleton to promote IT growth. For example, PIR1 (121F-specific p53 inducible RNA 1), NAP1 (Nck-associated protein 1), and SCARN (SCAR-Nodulation) (Yokota et al., 2009; Miyahara et al., 2010; Qiu et al., 2015) are components of an actin assembly SCAR/WAVE complex, and ARPC1 (actin related protein complex 1) is a predicted subunit of the actin-related protein complex ARP2/3 (Hossain et al., 2012). Another component is a legume-specific pectate lyase, NPL, which may be involved in cell wall remodeling during IT initiation (Xie et al., 2012). Rhizobia induce expression of root-hair-specific genes such as *CBS1* (Cystathionine-β-synthase-like 1), *RPG* (Rhizobium-directed polar growth), and *RINRK1* (Rhizobia infection receptor-like kinase 1) (Arrighi et al., 2008; Sinharoy et al., 2016; Li et al., 2019), although these genes have been identified but their biological functions in IT formation are not yet clear.

In response to rhizobia-secreted NFs, the root hair tips deform and entrap the bacteria; root hair cell nuclei then move to a location close to root hair tips and the plasma membrane invaginates to form an IT (Gage, 2004). Subsequent IT progression within root hairs follows the path of the moving nucleus, regardless of its direction, supporting the idea that nuclear movement is necessary for IT guidance (Fahraeus, 1957). The Linker of Nucleoskeleton and Cytoskeleton (LINC) complex in *M. truncatula* is necessary for proper nuclear shaping and movement in *Medicago* root hairs, and it plays a role in IT initiation and nodulation (Newman-Griffis et al., 2019). A cytoplasmic column, rich in secretory organelles, accumulates between the IT and the nucleus (Fournier et al., 2008) and encompasses a structure referred to as an infectosome (Liu et al., 2019b; Roy et al., 2020). In *L. japonicus*, the NF receptor NFR5 can interact with LjROP6 (Rho of Plants 6), which is activated by the DOCK family GEF (guanine nucleotide exchange factor) LjSPIKE1 (SPK1). LjSPK1-LjROP6 then guides polarized IT growth in root hairs (Ke et al., 2012; Liu et al., 2020). *L. japonicus* CERBERUS and its orthologue LIN (Lumpy infection) in *M. truncatula* displayed puncta localization and interacted with VAPYRIN, a protein of unknown function, to mediate IT polar growth (Murray et al., 2011; Liu et al., 2019b; Liu et al., 2021). Exo70H4, an exocyst subunit, co-localized with VAPYRIN and LIN during rhizobia infection, suggested that LIN, VAPYRIN and Exo70H4 may form a symbiosis-specific machinery to regulate polar growth of IT (Liu et al., 2019b).

NIN (Nodule inception) and RPG had been identified as two key genes which were lost in most non-nodulating species but essential for root nodule symbioses in nitrogen-fixing root nodule (NFN) clades including Fabales, Fagales, Cucurbitales, Rosales and Parasponia (Griesmann et al., 2018; van Velzen et al., 2018). A *M. truncatula rpg* mutant formed abnormally thick and slow-growing ITs, indicating that RPG plays an important role in IT tip growth (Arrighi et al., 2008). However, RPG has not been characterized in other legumes, and its precise biological function has remained elusive. In this study, we identified the *RPG* gene in *L. japonicus* and showed that it was required for IT formation. *RPG* displayed an infection-specific expression pattern and was directly induced by NIN. RPG showed punctate subcellular localization, could co-localize and interact with CERBERUS close to the nuclei in root hairs after rhizobia inoculation. We propose that the IT elongation, driven by movement of nucleus in root hairs is mediated by this infection-associated complex.

## RESULTS

### Identification of the *L. japonicus RPG* gene in infection-deficient mutants

Two symbiosis-defective mutants (SL5706-3 and SL454-2) were isolated from an ethyl methanesulfonate (EMS) mutagenized population in *L. japonicus* Gifu B-129. Both mutant lines produced small white nodules three weeks after inoculation (Figure 1, A-B and Supplemental Figure S1). SL5706-3 and SL454-2 were then crossed with the ecotype MG20 (Miyakojima) to generate mapping populations. F_1_ plants produced pink nodules two weeks after inoculation with *Mesorhizobium loti*. The nodulation phenotype was scored in F_2_ seedlings and revealed segregation of a monogenic recessive mutation in each line (SL5706-3: 145 nod^+^ and 37 nod^-^, χ^2^ value = 1.132; SL454-2: 237 nod^+^ and 60 nod^-^, χ^2^ value = 1.954). Rough mapping using established DNA markers (http://www.kazusa.or.jp/lotus/) revealed that both mutations were on *L. japonicus* linkage group 5, between markers TM0913 and TM0052. Two infection-related genes, *CERBERUS/LIN* and *RPG*, had previously been mapped to the corresponding region in *M. truncatula* (Arrighi et al., 2008; Kiss et al., 2009; Yano et al., 2009). We amplified and sequenced the genomic DNA corresponding to the coding region of *CERBERUS* and *RPG* in SL5706-3 and SL454-2. This revealed that neither line had a mutation in the *CERBERUS* gene, but each had a single point mutation in the putative ortholog of *RPG* (Lj5g3v1699100.1). SL5706-3 had a G to A transition at +3395 bp from the predicted start codon and SL454-2 had a G to A transition at +3650 bp; these mutations caused premature stops at residues W258 and W343 (Figure 1, C-D and Table 1).

**Figure 1.**
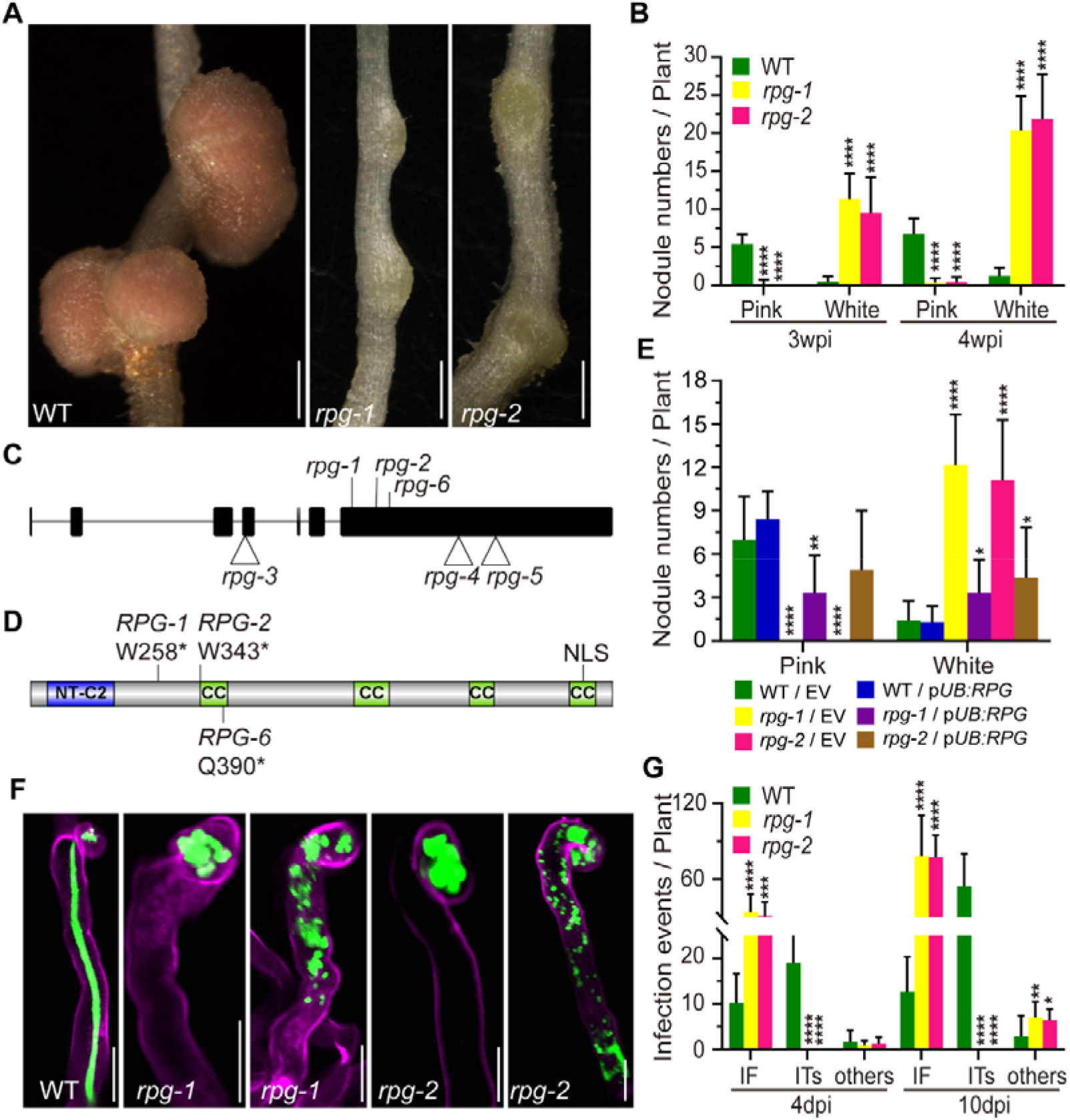
Phenotype and genotype of *rpg-1* and *rpg-2* mutants. A and B, Nodule phenotype (A) and the nodule numbers (B) of wild type (WT) and *rpg* mutants two weeks after inoculation. The total number of nodules per plant was scored at three and four weeks after inoculation with *M. loti* R7A LacZ (n>10). C, The gene structure of *RPG*, showing seven exons and six introns. The position of three EMS mutations, *rpg-1* (SL5706-3), *rpg-2* (SL454-2) and *rpg-6* (SL0181), and three *LORE1* insertion mutations, *rpg-3* (30053003), *rpg-4* (30055099), and *rpg-5* (30010526) are shown. D, Outline of RPG protein structure, indicating the predicted NT-C2 domain, four coiled-coil (CC) domains, and a predicted nuclear localization signal (NLS). The locations of translation stops in *RPG-1, RPG-2* and *RPG-6* are indicated. E, The nodule numbers of WT, *rpg-1* and *rpg-2* mutant plants in roots transformed with the vector control (EV) or p*UB:RPG* and scored three weeks after inoculation with *M. loti* R7A/LacZ (n>7). F, Infection phenotypes of WT, *rpg-1 and rpg-2* mutants were visualized by fluorescence microscopy of roots inoculated with *M. loti* R7A/GFP. Shown are a normal elongating IT in WT, and infection foci (IF) and abnormal ITs observed in *rpg-1 and rpg-2* mutants. Roots were scored seven days after inoculation and were counterstained with propidium iodide. Green fluorescence shows rhizobia and magenta fluorescence shows root hair stained with propidium iodide. G, Number of infection events in WT plants and *rpg* mutants. The total number of infection events per plant was scored 4 and 10 days after inoculation with *M. loti* R7A/LacZ. IF, infection foci; ITs, infection threads; Others correspond to abnormal ITs in root hairs as illustrated in panel F (n>9). Statistical significance was evaluated, comparison with WT. Scale bars: 1 mm (A); 20 μm (F).

**Table 1.**
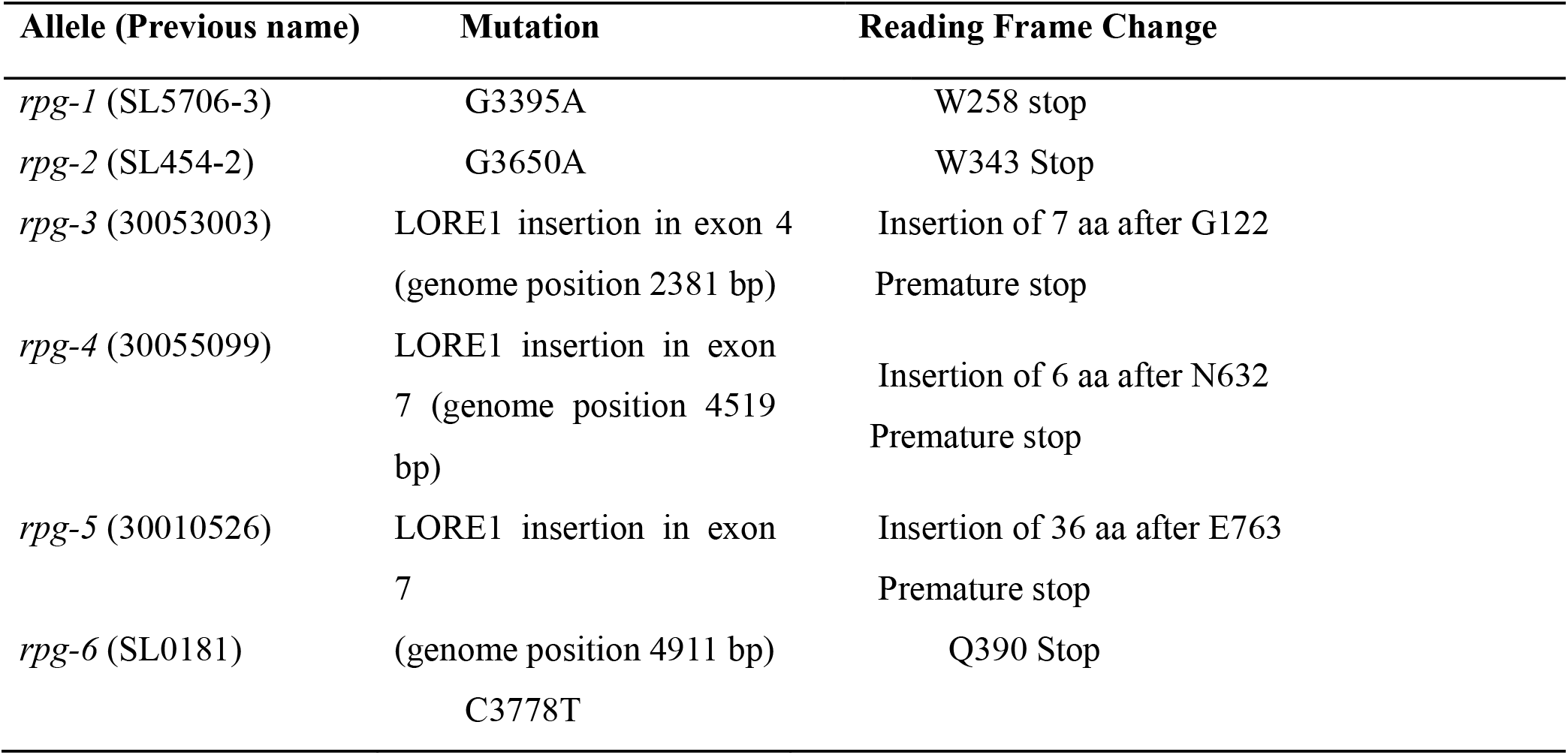
*L. japonicus rpg* mutant alleles.

| Allele (Previous name) | Mutation | Reading Frame Change |
| --- | --- | --- |
| *rpg-1* (SL5706-3) | G3395A | W258 stop |
| *rpg-2* (SL454-2) | G3650A | W343 Stop |
| *rpg-3* (30053003) | LORE1 insertion in exon 4<br>(genome position 2381 bp) | Insertion of 7 aa after G122<br>Premature stop |
| *rpg-4* (30055099) | LORE1 insertion in exon 7<br>(genome position 4519 bp) | Insertion of 6 aa after N632<br>Premature stop |
| *rpg-5* (30010526) | LORE1 insertion in exon 7 | Insertion of 36 aa after E763<br>Premature stop |
| *rpg-6* (SL0181) | (genome position 4911 bp)<br>C3778T | Q390 Stop |

To test if these mutations caused infection defects in the two mutant lines, wild type (WT) cDNA was amplified from Gifu mRNA and inserted into a plasmid under the control of the *L. japonicus* ubiquitin promoter. This construct (p*UB:RPG*) was introduced into SL5706-3 and SL454-2 by *Agrobacterium rhizogenes*-mediated hairy-root transformation, restoring normal nodulation in both mutants (Figure 1, E and Supplemental Figure S2). We conclude that the identified mutations in *RPG* caused the nodulation defect, and the alleles in SL5706-3 and SL454-2 were designated *rpg-1* and *rpg-2*, respectively.

*RPG* in *L. japonicus* is a 6.2-kb gene composed of seven exons separated by six introns (Figure 1, C). Reverse transcription and DNA sequencing indicated that the *LjRPG* cDNA is 3528 bp, encoding a protein 1176 amino acids in length. The predicted protein was 60% identical to MtRPG. The LjRPG protein domain was predicted to have four long coiled-coil domains, similar to MtRPG (Arrighi et al., 2008). As predicted for the *Parasponia* RPG (Zhang and Aravind, 2010; van Velzen et al., 2018), the LjRPG N-terminus had a predicted C2 (NT-C2) domain which predicted to mediate lipid-binding (Figure 1D) (Zhang and Aravind, 2010).

### Mutation of *RPG* blocks IT formation but not induction of early nodulation genes

Infection and nodulation phenotypes of the *rpg-1* and *rpg-2* mutants were analyzed after inoculation with *M. loti* R7A containing either a constitutively-expressed green fluorescent protein (GFP) or β-galactosidase (*lacZ*) marker gene. The WT plants produced elongated infection threads four days after inoculation (Figure 1, F), but most infection events in the *rpg* mutants were blocked at the stage of formation of infection foci (Figure 1, F). As observed with other infection-defective mutants (Murray et al., 2007; Yokota et al., 2009; Qiu et al., 2015; Li et al., 2019), bacteria were sometimes seen in the root hair cells (here we designated them as “others”) (Figure 1, F). Analysis of infection events in the *rpg* mutants four and ten days post inoculation (dpi) revealed that most infection events were arrested as infection foci, and neither mutant formed any normal-looking infections until 10 dpi (Figure 1, G).

Three *LORE1* insertion mutants (Fukai et al., 2012; Urbański et al., 2012) were obtained for *rpg* and the alleles were designated *rpg-3, rpg-4*, and *rpg-5* (Figure 1, C and Table 1). A third EMS-induced mutant line with an infection defect (SL0181) was also isolated and the mutation mapped in a similar manner to *rpg-1* and *rpg-2* (Supplemental Figure S3, A-B). SL0181 was found to have a C3778T transition in the sequence of *RPG* leading to a premature stop codon at Q390, then it was designated *rpg-6* (Figure 1, C-D and Supplemental Figure S3, C). Similar to the *rpg-1* and *rpg-2* mutants, most rhizobial infections in the *LORE1* insertion mutants did not go further than infection foci, although some infection threads were observed (Supplemental Figure S4, A-C). However, the *LORE1 rpg* mutants formed pink nodules three weeks after inoculation. The *rpg-3* mutant produced a similar number of mature-looking pink nodules as the WT, whereas *rpg-4* and *rpg-5* had fewer pink nodules (Supplemental Figure S4, B, D and E). The *rpg-6* mutant had a strongly reduced number of nodules and a high number of uninfected nodule primordia (Supplemental Figure S5, A-B). *RPG* expression was measured by quantitative reverse transcription (qRT)-PCR in all five *rpg* mutants. *RPG* transcript levels were significantly decreased in *rpg-1* and *rpg-2* but not in *rpg-3* or *rpg-4* mutants (Supplemental Figure S6).

*NIN, NPL, RINRK1* and *VPY1* are all induced by rhizobial infection (Schauser et al., 1999; Xie et al., 2012; Li et al., 2019; Liu et al., 2021). These genes were all expressed at similar levels in the *rpg-1* and *rpg-2* mutants as in the WT (Supplemental Figure S7) indicating that *RPG* is not required for the induction of rhizobial infection-related genes.

Arbuscular mycorrhization by *Rhizofagus irregularis* was also scored in the *rpg-1* and *rpg-2* mutants; microscopic examination and quantification of infections five weeks after inoculation identified no difference in hyphal penetration, or arbuscule formation compared with WT (Supplemental Figure S8) indicating that *RPG* is required for infection by rhizobia but not by arbuscular mycorrhizal fungi (AMF).

### *RPG* is induced by NIN and shows infection-specific expression

*RPG* transcript levels were increased in roots at several time points after inoculation with *M. loti* or after addition of purified *M. loti* NF (Figure 2, A-B). To investigate the spatial and temporal expression pattern of *RPG* during infection and nodulation, we used *A. rhizogenes* to transform *L. japonicus* WT roots with p*RPG:GUS*, which carries the *β-glucuronidase* (GUS) gene behind the *RPG* promoter. Expression could be detected in some epidermal cells (including root hairs) three to five days after inoculation with *M. loti* (Figure 2, C). We observed strong p*RPG:GUS* expression in infected root hairs; GUS staining co-localized with GFP-marked *M. loti* (Figure 2,D). Strong GUS staining was observed in nodule primordia, but there was much less staining in mature nodules (Figure 2,E-F). Sections of developing nodule primordia (at 5 dpi) revealed GUS expression in all cell layers (Figure 2,G), although GUS expression was then restricted to the nodule parenchyma cells in young mature nodules (14 dpi) (Figure 2, H).

**Figure 2.**
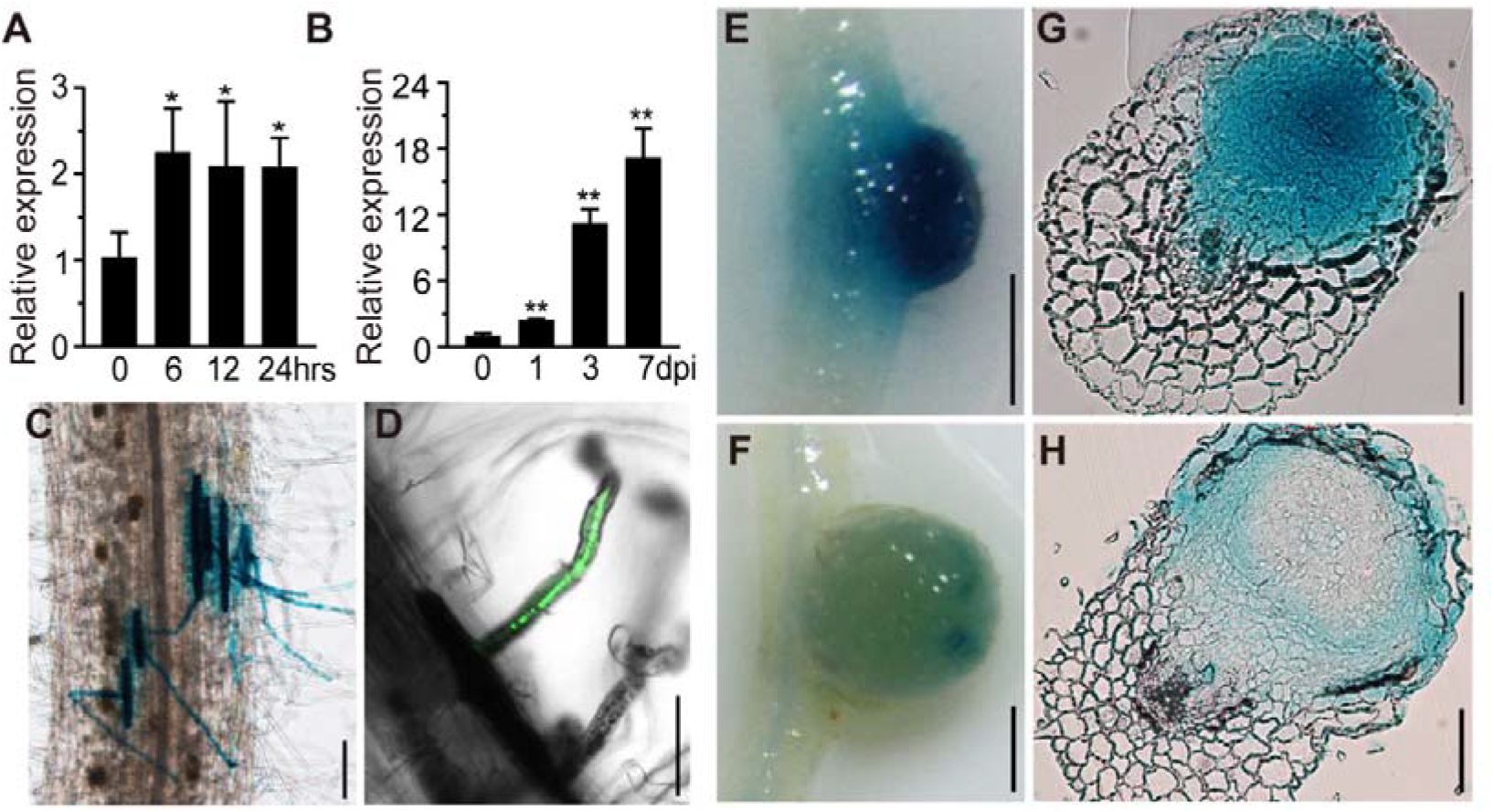
*RPG* expression pattern in *L. japonicus* roots. A and B, qRT-PCR analysis of *RPG* transcript levels in roots of wild type (WT) *L. japonicus*. Samples were collected at 0, 6, 12, and 24 h after inoculation with purified Nod factor (A) or at 0, 1, 3 and 7 days after inoculation with *M. loti* R7A (B). Expression is relative to that of mock-treated samples (0 h or 0 dpi) and normalized to *L. japonicus Ubiquitin*. Statistical significance was evaluated, comparison with 0 and indicated time points. C to F, p*RPG:GUS* expression in WT roots after inoculation with *M. loti* R7A (C, E, F) or *M. loti* R7A GFP (D). Strong GUS staining was detected in epidermal cells (C) and (D) and young nodules (E), but there was much lower staining in mature nodules (F). Bacteria are colored green to indicated infection threads (D). G and H, Nodule sections showed that *pRPG:GUS* induced GUS expression in all cell layers of young nodules (G), but GUS was only expressed in epidermal and nodule parenchyma cells in mature nodules (H). Scale bars: 100 µm (C-D and G-H); 1 mm (E-F).

To analyze how *RPG* expression is regulated by NF signaling, we measured *RPG* expression in *nin-2* and *ern1-2* mutants. This revealed that *M. loti*-induced *RPG* expression requires *NIN* and *ERN1* in *L. japonicus* roots (Figure 3, A). We then used a dual-luciferase (dual-LUC) reporter assay to analyze whether NIN or ERN1 directly affects *RPG* transcription by co-expressing p*RPG:LUC* with p*35S:NIN* or p*35S:ERN1* in *Nicotiana benthamiana* leaf cells. Luciferase activity was quantified in leaf discs, revealing that NIN, but not ERN1, could induce *RPG* expression (Figure 3, B). Two putative NIN-binding nucleotide sequences (NBS) (Soyano et al., 2014) were identified 1157 bp (S1) and 241 bp (S2) upstream of the *RPG* translation start codon (Supplemental Figure S9). We used an electrophoresis mobility shift assay (EMSA) to determine whether NIN could bind to these regions of the *RPG* promoter. A mobility shift was observed when the carboxyl-terminal half of the NIN recombinant protein was incubated with a synthetic oligonucleotide corresponding to the identified sequence in the *RPG* promoter S1 and S2 regions; an unlabeled competitor oligonucleotide outcompeted binding by the labelled probe (Figure 3, C-D). Deletion of the conserved NBS of the S2 region (ΔS2, -29 bp) prevented NIN binding, but deletion of the S1 region (ΔS1, -35 bp) did not blocked NIN binding (Figure 3, C-D). In a competition assay, unlabeled ΔS2 could not outcompete NIN binding to the labelled S2 region, whereas unlabeled ΔS1 could outcompete the NIN binding to the S1 probe (Figure 3, C-D). These results all suggest that the S2 region is critical for NIN binding to the *RPG* promoter. To verify this, we used a dual-LUC system with p*RPG:LUC* containing deletions of S1 (p*RPGΔS1:LUC*), S2 (p*RPGΔS2:LUC*) or with both S1 and S2 deleted (p*RPG Δ S1,2:LUC*) and co-expressed each with p*35S:NIN* in *N. benthamiana* leaves. The results showed that NIN could not induce expression of p*RPGΔS2:LUC* or p*RPGΔS1,2:LUC*, but could induce p*RPGΔS1:LUC* expression (Figure 3,B). This indicates that S2 is essential for induction of *RPG* by NIN. The results were validated in *L. japonicus* by expressing p*RPG*:*GUS*, p*RPGΔ S1:GUS*, or p*RPGΔS2:GUS* in transformed *L. japonicus* hairy roots; p*RPG:GUS* and p*RPG*ΔS1*:GUS* had similar expression patterns in roots inoculated with *M. loti* (Figure 3, E-F). In contrast, about half of the p*RPG*Δ*S2:GUS* transgenic roots (13/27) had no detectable GUS expression (Figure 3, H) and the remainder (14/27) showed weaker GUS staining than *pRPG:GUS* (Figure 3,G). Based on these observations, we conclude that *RPG* is induced by NIN through an interaction with the S2 region, resulting in an infection-specific expression pattern.

**Figure 3.**
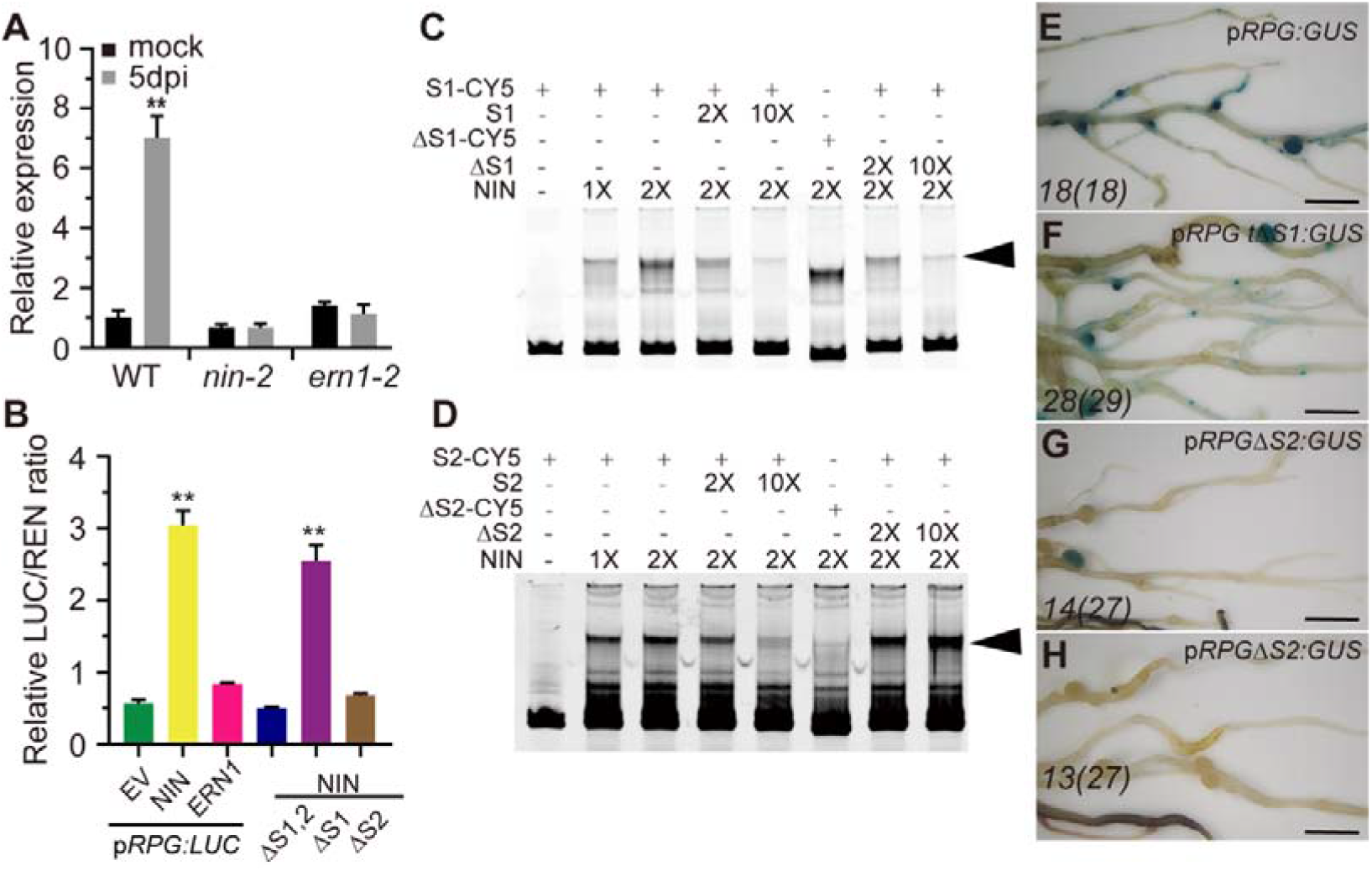
NIN induces *RPG* expression. A, qRT-PCR analysis of *RPG* transcript levels in WT, *nin-2*, or *ern1-2* roots 5 days after inoculation with *M. loti* R7A. Expression is relative to that of mock-inoculated WT and normalized to *L. japonicus Ubiquitin*. Statistical significance was evaluated, comparison with mock-inoculated WT. B, Transcriptional activation ability of ERN1 or NIN to *RPG* promoter in *N. benthamiana* leaves. Co-expressed effector (ERN1 or NIN) and a double-reporter plasmid contain RPG promoter driven LUC reporter gene and Renilla Luciferase (REN) driven by CaMV35S in *N. benthamiana* leaves. *Renilla* Luciferase (REN) activity was used to normalize for the efficiency of transformation. Statistical significance was evaluated, comparison with empty vector (EV). C and D, Gel-shift assays of NIN binding to the promoter of *RPG*. DNA fragments (1 nM) from portions of the *RPG* promoter carrying the S1 and S2 regions were amplified by PCR, fluorescently labeled (CY5), and incubated with the indicated concentrations of NIN protein for 20 min at 30 °C. The protein-DNA complexes were separated by electrophoresis on native 6% polyacrylamide gels, and the fluorescently-labelled DNA was detected by fluorimetry. For S1 and S2, a 2-fold and 10-fold excess of unlabeled DNA fragments were added as competitors for binding. E to H, Assays of p*RPG:GUS*, p*RPGΔS1:GUS*, p*RPGΔS2:GUS* expression in transformed *L. japonicus* roots. GUS activity was similar in p*RPG:GUS* (E) and p*RPGΔS1:GUS* (F), but showed reduced levels in p*RPGΔS2:GUS* (G) and (H). Scale bars: 5 mm (E-H).

### RPG displays punctate subcellular localization

To investigate the subcellular localization of RPG, we first used assays in *N. benthamiana* leaves. We had expected RPG to be localized to the nucleus based on prior results with *Medicago* RPG (Arrighi et al., 2008) and the predicted nuclear localization signal (NLS) (http://www.psort.org/) at the C-terminus of LjRPG (Figure 1,D). However, GFP-RPG made by fusing *GFP* with *RPG* cDNA and expressed by the 35S promoter showed strong fluorescence with punctate foci, some of which were close to the nucleus (Figure 4, B). Expressing the p*35S:GFP-RPG* construct in the *rpg-1* mutant can rescue its infection defects and produce pink mature nodules (Supplemental Figure S10). This observation was confirmed in *L. japonicus* root protoplasts in which *L. japonicus* ASTRAY, a homologue of *Arabidopsis thaliana* HY5 (Nishimura et al., 2002), was used as a nuclear marker. Co-expression of GFP-RPG and ASTRAY-mRFP in *L. japonicus* root protoplasts revealed puncta of GFP-RPG, some of which were close to, but distinct from the nucleoplasm (Figure 4, D). As expected, expression of GFP alone (p*35S:GFP*) showed both nuclear and cytoplasmic localization (Figure 4,A and C).

**Figure 4.**
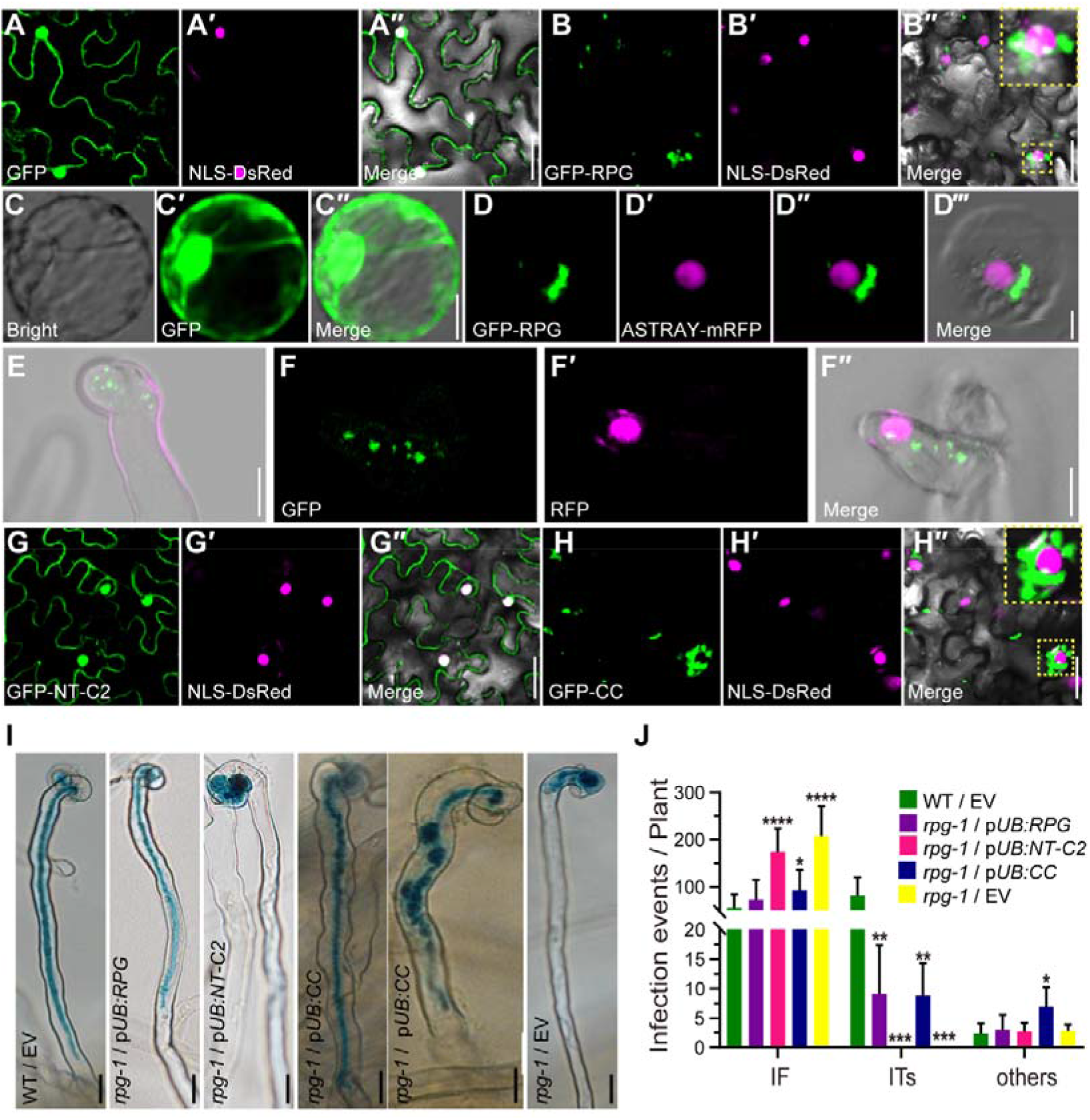
Subcellular localization of the RPG protein. A, B, G and H, Confocal microscopy images of *N. benthamiana* leaves expressing p*35S:GFP* (A), p*35S:GFP-RPG* (B), and the separate RPG domains RPG p*35S:GFP-NT-C2* (G) and p*35S:GFP-CC* (H). In each, the green, magenta and merged images are shown in adjacent panels. The nucleus was labeled with NLS-DsRed (magenta). Sections within an image that are outlined in dotted yellow lines are showed enlarged in the top right corner of that image. C and D, p*35S:GFP* (C) or p*35S:GFP-RPG* (green) and the nuclear marker, ASTRAY5-mRFP (magenta) (D), were co-expressed in *L. japonicus* root protoplasts using a DNA-PEG-calcium transfection method. E and F, p*RPG:GFP-RPG* was introduced into *rpg-1* by *A. tumefaciens*-mediated stable transformation. To detect GFP-RPG subcellular localization 5 days after inoculation with *M. loti* MAFF303099/RFP, the roots were analyzed using whole-mount immunolocalization with GFP primary antibody and Alexa Fluor 488-conjugated Affinipure donkey anti-Mouse IgG secondary antibody. Green shows RPG subcellular localization and magenta shows *M. loti*. I and J, Assays of complementation of the *rpg-1* mutant by the predicted NT-C2 domain (p*UB:NT-C2*) and by RPG lacking the NT-C2 domain (p*UB:CC*) showing the NT-C2 domain is not required for complementation of infection. Infection phenotypes (H) and Infection events (I) of the RPG NT-C2 or CC domain were driven by p*UB* promoter and expressed in *rpg-1* using hairy root transformation after inoculation with *M. loti* R7A/LacZ (n>14). Statistical significance was evaluated, comparison with empty vector WT/EV. Scale bars: 25 µm (A-B and G-H); 10 µm (C-F); 20 μm (I).

To analyze RPG subcellular localization in legumes after rhizobial inoculation, the *L. japonicus rpg-1* mutant was stably transformed with *GFP* fused to *RPG* cDNA downstream of the native *RPG* promoter (p*RPG:GFP-RPG*). Analysis of T_2_ plants of this transformant revealed that those that expression of p*RPG:GFP-RPG* in the *rpg-1* mutant resulted in formation of normal ITs and pink nodules as seen in the WT. In contrast, T_2_ segregants lacking p*RPG:GFP-RPG* (*rpg-1*) formed infection foci and white nodules as seen in the mutant (Supplemental Figure S11). This shows that the GFP-RPG fusion protein functioned in the transgenic roots of the *rpg-1* mutant. No GFP fluorescence could be reliably detected in live roots, so we immuno-localized the protein using GFP antiserum. There was little or no detectable signal in the absence of *M. loti*, but punctate localization of GFP-RPG was observed in root hairs with infection foci after inoculation with *M. loti* MAFF303099/RFP (Figure 4, E-F) or ITs (Supplemental Figure S12, A), and was also observed in inoculated root hairs that did not contain entrapped rhizobia (Supplemental Figure S12, B). Taken together, these results showed that RPG localizes in puncta close to the nucleus when expressed in *N. benthamiana* leaves or *L. japonicus* root protoplasts, also showed puncta dots in *L. japonicus* root hairs following inoculation with *M. loti*.

To analyze the domain of RPG that determines its subcellular localization, we made constructs in which the GFP was fused either to the RPG N-terminal NT-C2 domain contained in the first 300 amino acids of RPG (GFP-NT-C2) or the region of the protein (residues 170-1176) lacking the NT-C2 domain but containing all the C-terminal coiled-coil domains (GFP-CC). In *N. benthamiana* leaves GFP-NT-C2 was expressed throughout cells (Figure 4, G), similar to free GFP (Figure 4, A), whereas GFP-CC displayed the same punctate localization as full-length GFP-RPG (Figure 4, B and H); protein levels were quantified by immunoblotting with anti-GFP antiserum (Supplemental Figure S13). The observed localization suggested that the NT-C2 domain is not required for the observed subcellular localization of RPG. Constructs were then generated in which the NT-C2 or the protein lacking the NT-C2 domain were expressed by the *L. japonicus Ubiquitin* promoter (Maekawa et al., 2008); these were expressed in roots of *rpg-1* using hairy root transformation. Expression of the *RPG* lacking the NT-C2 domain (p*UB:CC*) rescued the *rpg-1* infection defect as effectively as full-length *RPG* and the transformants formed normal ITs and pink mature nodules (Figure 4, I-J and Supplemental Figure S14). No rescue was observed in roots expressing NT-C2 (p*UB:NT-C2*) (Figure 4, I-J and Supplemental Figure S14) and in all cases transformation was confirmed with a separate GFP marker. Based on these data, we conclude that RPG displays punctate subcellular localization, and that the N-terminal C2 domain is not required for this subcellular localization or its biological function.

### RPG interacts with CERBERUS at the TGN/EE compartment

The punctate localization of RPG is similar to that reported for MtLIN and LjCERBERUS (Liu et al., 2019b; Liu et al., 2021). We hypothesized that RPG and CERBERUS may function together to promote IT formation. To define the subcellular compartment in which RPG localized, we co-expressed GFP-RPG or mCherry-RPG with the following subcellular markers in *N. benthamiana* leaves: Sec12-PHB-mCherry for the endoplasmic reticulum (ER); HAP3-GFP or SYP61-mCherry for the TGN/EE; ARA6-mCherry or mRFP-VSR2 for multivesicular bodies (MVB); and CD3-963-GFP for the Golgi (Bar-Peled and Raikhel, 1997; Ueda et al., 2001; Nelson et al., 2007; Drakakaki et al., 2012; Wang et al., 2013). The results showed that RPG co-localized with ER and TGN/EE markers (Figure 5, A and Supplemental Figure S15, A-B) but not with the Golgi or MVB markers (Supplemental Figure S15, C-E). This is similar to the observed subcellular localization of CERBERUS and indeed, co-expressed GFP-RPG and CERBERUS-mCherry in *N. benthamiana* leaves co-localized to punctate loci (Figure 5,B).

**Figure 5.**
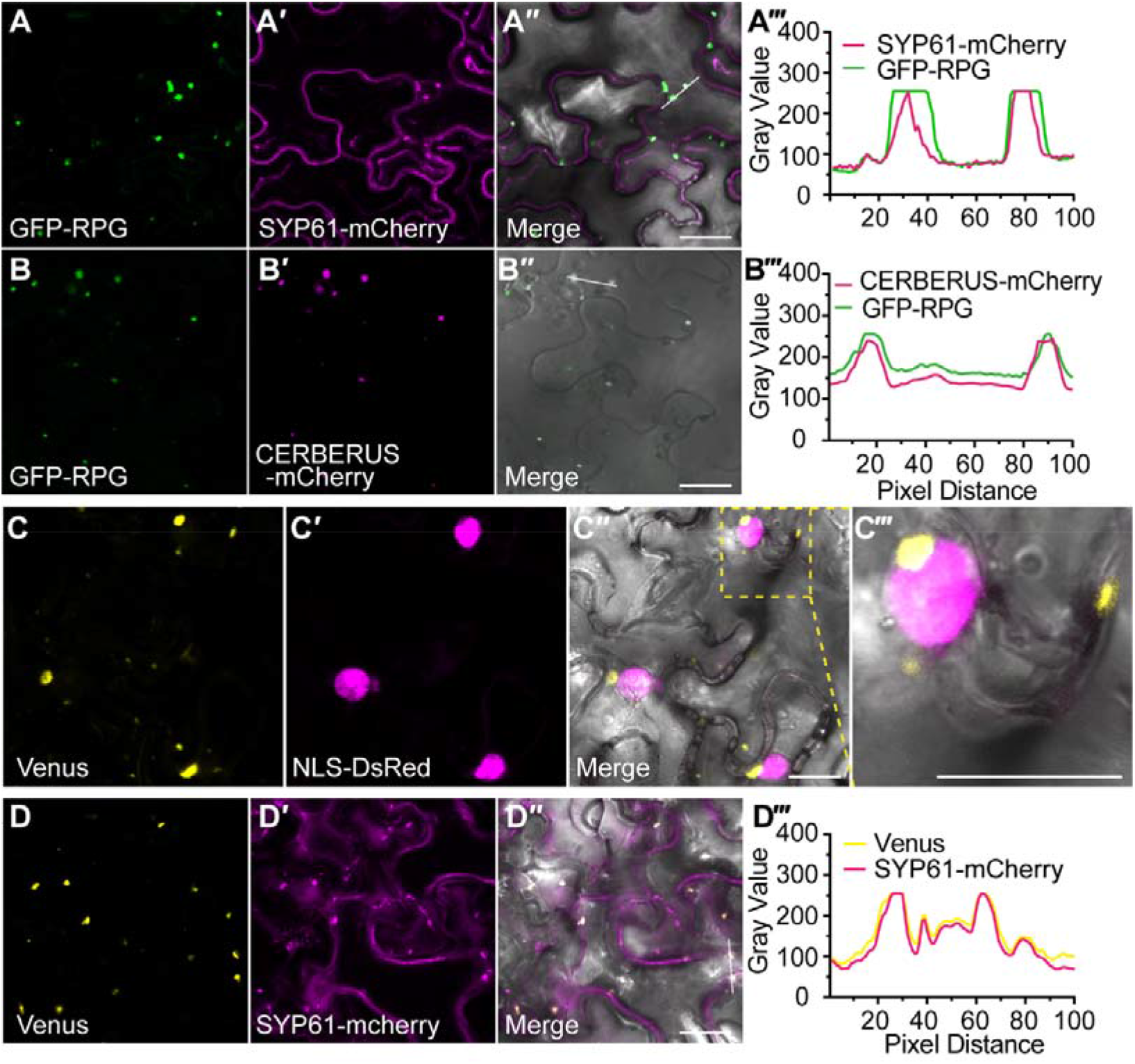
RPG co-localization with a TGN/EE marker and interaction with CERBERUS. A and B, GFP-RPG (green) and TGN/EE marker SYP61-mCherry (magenta) (A) or CERBERUS-mCherry (magenta) (B) were co-expressed in *N. benthamiana* leaf cells. The images show GFP, mCherry and merged fluorescence images. Plots (A‴) and (B‴) show fluorescence intensities of GFP-RPG and SYP61-mCherry or CERBERUS-mCherry in regions of interest (indicated by white line in [A″] and [B″]). C, Co-expressed RPG-CERBERUS BiFC construct (yellow) and NLS-DsRed (magenta) in *N. benthamiana* leaf cells. The image shows strong Venus fluorescence localized in puncta, some of which were close to the nucleus. (C‴) shows an enlargement of the area in outlined in yellow in the merged image (C″). D, Co-expressed RPG-CERBERUS BiFC construct (yellow) and TGN marker SYP61-mCherry (magenta). The image shows merging of Venus (yellow) and TGN marker fluorescence. Plot (D‴) shows fluorescence intensities of Venus and SYP61-mCherry in regions of interest (inset in [D″]). Scale bars: 25 µm (A-D).

We used a bimolecular fluorescence complementation (BiFC) assay in *N. benthamiana* using split-Venus fused to RPG and CERBERUS to determine if RPG can associate with CERBERUS. Co-expression of nVenus-CERBERUS and cVenus-RPG resulted in strong Venus fluorescence in puncta, some of which were close to the nucleus, which was marked with nuclear localized DsRed (NLS-DsRed) (Figure 5, C). Co-expression of a BiFC construct (containing p*AtUBI10*:*nVenus-CERBERUS* and p*LjUBI1*:*cVenus-RPG*) with markers for the ER (Sec12-PHB-mCherry) and TGN/EE (SYP61-mCherry) in *N. benthamiana* leaves confirmed that the RPG-CERBERUS complex co-localized with ER and TGN/EE markers (Figure 5, D and Supplemental Figure S15F), suggesting that the RPG–CERBERUS may interact to function in endosome trafficking during IT formation.

We used multiple assays to check which domains of RPG and CERBERUS mediate their interaction. Yeast two-hybrid (Y2H) assays confirmed that RPG could interact with full length CERBERUS, and showed that the CERBERUS Armadillo-like domain (ARM) (but not the WD40 domain) and the RPG coiled-coil domain (CC) were sufficient for the interaction (Figure 6, A). Split-luciferase complementation imaging assays in *N. benthamiana* leaves confirmed that CERBERUS could interact with RPG, and showed that the CERBERUS ARM domain interacted more strongly than full-length CERBERUS (Figure 6, B). Interaction of RPG and CERBERUS was also detected in a co-immunoprecipitation (Co-IP) assay in which GFP-RPG was co-expressed with CERBERUS-mCherry in *N. benthamiana* leaves (Figure 6, C).

**Figure 6.**
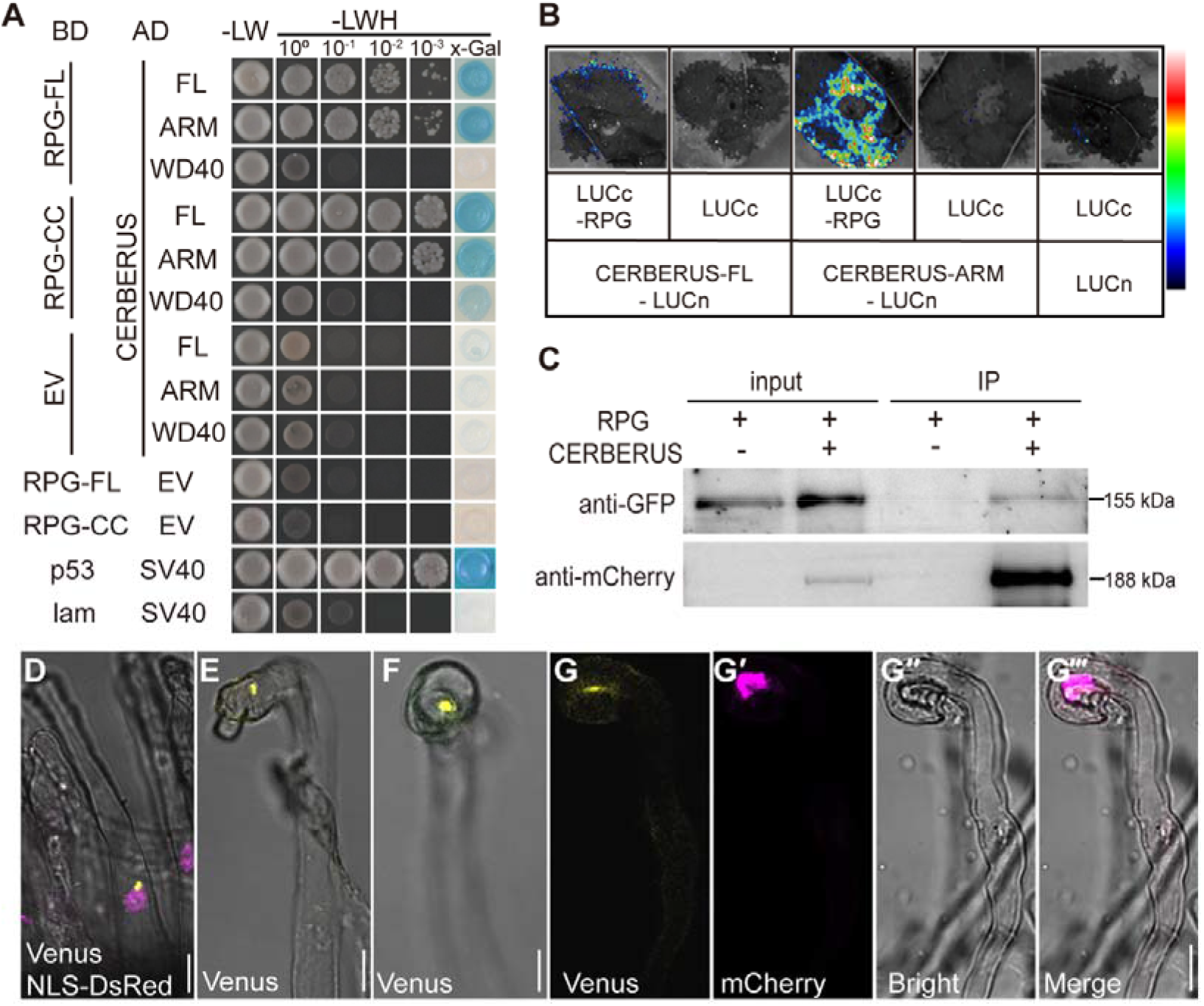
RPG interactions with CERBERUS. A, A GAL4-based yeast two-hybrid system was used to analyze the interaction between CERBERUS and full-length RPG (RPG-FL) or RPG lacking the NT-C2 domain (CC), and between CERBERUS ARM or WD40 and RPG-FL or RPG-CC. Potential interactions were assayed by growth on SD/-LWH (medium without histidine, leucine, or tryptophan) after gradient dilution. Images show the growth of co-transformants on selection media after three days. B, Luciferase biomolecular complementation assays of the interaction between RPG and full-length CERBERUS or CERBERUS ARM in *N. benthamiana* leaf cells. The indicated constructs were transiently co-expressed in *N. benthamiana* leaves, and luciferase complementation imaging was conducted two days after agroinfiltration. LUCn, N-terminal fragment of firefly luciferase. LUCc, C-terminal fragment of firefly luciferase. Fluorescence signal intensity is indicated. C, Co-Immunoprecipitation (Co-IP) assays of interactions between GFP-RPG and CERBERUS-mCherry in *N. benthamiana* leaves. GFP-RPG and CERBERUS-mCherry were co-expressed in *N. benthamiana* leaves. Co-IP was assayed using anti-mCherry antibody, and the precipitated proteins were detected by immunoblot analysis with anti-mCherry and anti-GFP antibodies. One representative results out of two biological replicates is shown. D to G, RPG-CERBERUS BiFC construct was expressed in *M. truncatula sunn-1* by hairy root transformation. To detect RPG and CERBERUS interaction, the transgenic roots were observed seven days after inoculation with Sm1021/mCherry. Venus Fluorescence (yellow) shows RPG and CERBERUS interaction. NLS-DsRed (magenta) indicates the nucleus (D). mCherry (magenta) shows rhizobium Sm1021/mCherry in (G′ and G‴). Scale bars: 10 µm (D-G).

We further validated the RPG–CERBERUS interaction using BiFC in legumes. Transformation of *L. japonicus* roots with p*AtUBI10:nVenus-CERBERUS* and *pLjUBI1:cVenus-RPG* was selected using NLS-DsRed as a marker of transformation but we could not reproducibly detect Venus fluorescence. To try to enhance sensitivity, the same BiFC construct was expressed in roots of the *M. truncatula sunn-1* mutant, which shows increased levels of gene expression due to lack of autoregulation of nodulation (Schnabel et al., 2010). After inoculation with *Sinorhizobium meliloti* 1021, which carried an mCherry reporter, punctate Venus fluorescence was detected in root hairs close to the nucleus (Figure 6, D) and in curled root hairs (Figure 6, E-F), and was co-localized with bacteria in the curled root hair (Figure 6, G). These puncta labelled with the interacting proteins were similar to those labelled by GFP-LIN in *M. truncatula* PITs (Liu et al., 2019b). Based on these results, we conclude that RPG interacts with CERBERUS in infected root hairs.

## DISCUSSION

RPG was identified as a key gene in nodulating FaFaCuRo clades (Griesmann et al., 2018; van Velzen et al., 2018), which was required for IT polar growth in *M. truncatula* (Arrighi et al., 2008), and play more important roles in root hair ITs than transcellular ITs in *L. japonicus* (Montiel et al., 2021). In this study, we showed in *L. japonicus rpg* showed similar symbiotic phenotype as other infection threads deficient (*itd*) mutants, such as *rinrk1, scarn, cerberus* and *npl* (Lombardo et al., 2006; Yano et al., 2009; Xie et al., 2012; Qiu et al., 2015; Li et al., 2019). However, *RINRK1* was required for rhizobia-induced nodulation genes expression (Li et al., 2019), while the induction of these genes by rhizobia was not affected in *scarn* and *rpg* mutants (Qiu et al., 2015 and this study), suggested that the biological roles of SCARN and RPG in IT formation were different with RINRK1, which may involve in NF signaling transduction pathway.

*RPG* is induced by either purified NF or inoculation with *M. loti*. It was previously reported that *MtRPG* expression is dependent on NIN (Liu et al., 2019a), and chromatin immunoprecipitation also showed that NIN could directly bind with the *RPG* promoter in *L. japonicus* roots (Soyano et al., 2014). We found that the induction of *RPG* by rhizobia requires ERN1 and NIN, whereas NIN but not ERN1, could directly bind to the *RPG* promoter to activate *RPG* expression. This may be because ERN1 functions upstream of NIN, as has been suggested in *M. truncatula* (Hirsch and Oldroyd, 2009). Thus, ERN1 induction of *NIN* could induce *RPG* expression. Two predicted NIN binding sites were present in the *RPG* promoter, and although NIN could directly bind to both *in vitro*, but, only one was required for *RPG* induction in roots.

RPG contains a predicted NT-C2 domain and four predicted long coiled-coil domains. Proteins containing the NT-C2 domain, such as vertebrate estrogen early-induced gene1 (EEIG1) (Wang et al., 2004) and its ortholog in *Drosophila*, are required for uptake of dsRNA *via* the endocytic machinery to induce RNAi silencing (Saleh et al., 2006). In *Arabidopsis*, PLASTID MOVEMENT IMPAIRED 1 (PMI1) is a plant-specific C2-domain protein that is required for efficient movement of chloroplasts nuclei in response to light (DeBlasio et al., 2005; Suetsugu et al., 2015). However, in this study, we did not observe a requirement for the predicted NT-C2 domain for IT formation suggesting that the lipid binding activity of RPG NT-C2 domain should be tested. RPG C-terminal long coiled-coil domain is sufficient for its protein subcellular localization and biological function. Long coiled-coils are highly versatile protein folding motif which function involved in organelle architecture, in nuclear organization and the cytoskeletal motor proteins (Rose et al., 2004; Truebestein and Leonard, 2016). MtRPG can interact with MtIEF (Infection-related epidermal factor), a legume-specific protein which contain a coiled-coil region and a DUF761 domain with unknown function (Kovács et al., 2022). In this study, we found that RPG was localized in puncta close to the nucleus in *N. benthamiana* leaves and *L. japonicus* root protoplasts. A very recent study in *M. truncatula* showed that the IT tip-to-nucleus microtubule connectivity is perturbed in *rpg-1* (Lace et al., 2022). Together all these studies strongly indicated that RPG plays important roles in nuclear-led IT elongation. MtLIN (the orthologue of CERBERUS) is localized at PIT tips, and also localizes in clear puncta associated with the nucleus (Liu et al., 2019b). RPG and CERBERUS interact near the nucleus and TGN/EE compartment; this suggests that the RPG–CERBERUS complex could promote polar growth of ITs by affecting nuclear migration through a connection to endosome trafficking and/or cytoskeletal changes during IT formation.

CERBERUS contains a U-box domain and has auto-ubiquitination activity (Yano et al., 2009; Liu et al., 2021). CERBERUS interacts with LjVPY1/2, but promotes LjVPY1/2 accumulation in *N. benthamiana* and *L. japonicus* (Liu et al., 2021). Despite persistent efforts, we were unable to express and purify RPG from *Escherichia coli*, and RPG expression in *N. benthamiana* was too low to perform an *in vitro* ubiquitination assay for CERBERUS and RPG. Moreover, LIN-VPY-Exo70H4 form a protein complex in infectosome in *M. truncatula* root hairs (Liu et al., 2019b). It will be very important to determine in the future the detailed molecular mechanisms of how the RPG, CERBERUS, VAPRYIN, and exocyst polar growth machinery operates in IT formation.

## MATERIALS and METHODS

### Plant materials and strains

The *L. japonicus* ecotypes Gifu B-129 and Myakojima (MG-20) and mutant lines *nin-2, ern1-2* (Cerri et al., 2017), and *cerberus-12* (Liu et al., 2021) were used in this study. For *M. truncatula*, the mutant line *sunn-1* (Schnabel et al., 2005) was used. The mutant lines *rpg-1* (SL5706-3), *rpg-2* (SL454-2), and *rpg-6* (SL0181) were isolated from forward genetic screening of an EMS mutagenesis population of *L. japonicus* Gifu B-129. Other *rpg* allelles were obtained from a *LORE1* retrotransposon insertion mutagenesis pool (Urbański et al., 2012). The transposon insertion in each gene was verified by PCR product sequencing; primers are shown in Supporting Information supplemental Table S1. *Meshorhizobium loti* R7A, constitutively expressing GFP or *lacZ* (referred to as R7A GFP or R7A LacZ), or *M. loti* MAFF303099 carrying RFP, or DsRED were used for *L. japonicus* nodulation experiments, and *Sinorhizobium meliloti* 1021-mCherry was used for *M. truncatula* nodulation experiments. Spores of the mycorrhizal fungus *Rhizophagus irregularis* were used for analysis of AM symbiotic phenotypes. For hairy root transformation of *L. japonicus* or *M. truncatula* roots, *Agrobacterium rhizogenes* strain AR1193 was used. *A. tumefaciens* strain EHA105 or GV3101 (pSoup) were used for *N. benthamiana* transient expression and stable transformation of *L. japonicus* as previously described (Tirichine et al., 2005). Plasmids were cloned in *Escherichia coli* DH10B or DH5α and *E. coli* Rosetta was used for protein expression. *Saccharomyces cerevisiae* strain AH109 was used for the yeast two-hybrid assay.

### Cloning, DNA manipulation, and plasmid construction

For genetic complementation, the coding sequence (CDS) of RPG was amplified from a cDNA library of inoculated Gifu roots using the primers *RPG-XbaI-F* and *RPG-AscI-R*. The PCR products and pUB-GFP plasmid were digested with *XbaI* and *AscI*, then RPG was inserted into pUB-GFP to form p*UB:RPG*. The NT-C2 and entire CC domains of RPG were amplified by PCR using the primers *RPG-attB-F/NT-C2-attB-R* or *CC-attB-F/RPG-attB-R*. The PCR product was inserted into pDONR207 via a BP reaction (Invitrogen, Waltham, MA, USA) and combined into pUB-GW-GFP to generate the p*UB:NT-C2* or p*UB:CC* construct via the LR reaction (Invitrogen).

For yeast two-hybrid assays, *RPG* PCR products were recombined into pDONR207 via a BP reaction. RPG/pDONR207, NT-C2/pDONR207, and CC/pDONR207 were recombined into pDEST-GBKT7 or pDEST-GADT7 using the LR reaction.

For split-luciferase complementation imaging assays, *RPG, CERBERUS*, and a fragment encoding the CERBERUS ARM domain were inserted into the destination vectors 771-LUCn and 772-LUCc following *KpnI* and *SalI* digestion.

To determine the subcellular localization of RPG in *N. benthamiana* leaves, RPG/pDONR207, NT-C2/pDONR207, and CC/pDONR207 were recombined into destination vector pK7WGF2-NLS-DsRed, which was modified from pK7WGF2. The kanamycin resistance gene of pK7WGF2 was replaced with a fragment of *NLS-DsRed* which was driven by the ubiquitin promoter. To obtain stably transformed plants, RPG/pDONR207 was recombined into pK7WGF2. The *RPG* promoter was amplified to replace the 35S promoter in RPG/pK7WGF2 to generate p*RPG:GFP-RPG*.

For co-localization and BiFC analyses in *N. benthamiana* leaves, *L. japonicus* plants, and *M. truncatula* hairy roots, mCherry-RPG constructs (co-localization assays) or RPG and CERBERUS constructs (BiFC assays) were generated with Golden Gate cloning (Weber et al., 2011). The RPG and CERBERUS CDS were synthesized in the level 0 vector pL0V-SC3 (Shanghai Xitubio Biotechnology) to generate p*L0M-SC3-RPG* or p*L0M-SC3-CERBERUS*. p*L0M-SC3-RPG* and the EC15111 vector were digested with BsaI to generate mCherry-RPG as the level 1 construct. This level 1 mCherry-RPG was assembled into EC50507 (https://www.ensa.ac.uk/) to generate the level 2 construct mCherry-RPG binary vector. p*L0M-SC3-RPG* was assembled into EC10048 to generate cVenus-RPG, and p*L0M-SC3-CERBERUS* was assembled into EC10044 to generate nVenus-CERBERUS. Finally, these constructs were assembled into EC50507, adding p*35S:NLS-DsRed* or p*35S:GUS* as a transgenic marker, to generate the BiFC construct p*AtUBI10:nVenus-CERBERUS/*p*LjUBI1:cVenus-RPG*. For transient expression in *L. japonicus* root protoplasts, the *GFP-RPG* fragment was amplified using RPG/pK7WGF2-NLS-DsRed as a template. The PCR products and pA7-GFP were digested with SpeI and BamHI, then GFP-RPG was inserted into pA7-GFP to form pA7-GFP-RPG. *ASTRAY* cDNA was amplified from Gifu cDNA using the primers *ASTRAY-BamHI-F* /*ASTRAY-BamHI-R*. The PCR products and pA7-mRPF were digested with BamHI, then inserted into pA7-mRFP to generate ASTRAY-mRFP by homologous recombination methods (Vazyme).

For the dual-luciferase reporter assay in *N. benthamiana*, the *RPG* promoter fragments or those with NBSs deleted were amplified via PCR using the primers shown in Supplemental Table S1. Single or multiple PCR products were then inserted into the pGreen II vector using homologous recombination methods (Vazyme) following KpnI and HindIII/SpeI digestion to generate the p*RPG:LUC*, p*RPGΔS1:LUC*, p*RPGΔS2:LUC*, and p*RPGΔS1,2:LUC* constructs. The effector construct was generated by inserting NIN or ERN1 CDS into the pRI101 vector (containing the 35S promoter) using the KpnI/EcoRI restriction sites.

For expression analysis of RPG in *L. japonicus* hairy roots, *RPG* promoter fragments (1581 bp upstream of their respective start codons) were amplified from genomic DNA extracted from Gifu leaves by PCR. The *RPG* fragments with NBS S1 or S2 deleted from the promoter were amplified using p*RPGΔS1:LUC* or p*RPGΔS2:LUC* as a template. PCR products were cloned into pDONR207 with a BP reaction, and combined into pKGWFS7-NLS-DsRed to generate the p*RPG:GUS*, p*RPGΔS1:GUS* or p*RPGΔS2:GUS* constructs via LR reaction. The pKGWFS7.0-NLS-DsRed vector was modified from pKGWFS7.0 by replacing the kanamycin with the *ubiquitin drive NLS-DsRed*.

All PCR amplification was performed using MAX (Vazyme), and all constructs were confirmed by DNA sequencing. Primers are shown in Supplemental Table S1 and constructs are listed in Supplemental Table S2.

### Map-based cloning

The *rpg-1, rpg-2* and *rpg-6* mutations were mapped using an F_2_ populations established by using SL5706-3, SL454-2 and SL0181 as pollen donors to *L. japonicus* ecotype MG20. Plants were inoculated with *M. loti* R7A LacZ and scored at 21 dpi for the nodulation phenotype. Genomic DNA was extracted from leaves as previously described (Pajuelo and Stougaard, 2005). Primer sequences and information for SSR markers were retrieved from the miyakogusa.jp website (http://www.kazusa.or.jp/lotus/).

### Plant growth conditions, symbiotic inoculations, and phenotype observation

*L. japonicus* or *M. truncatula* seeds were scarified, surface sterilized, and grown as previously described (Qiu et al., 2015; Luo et al., 2021). After five to seven days of growth, seedlings were inoculated with *M. loti* R7A LacZ, MAFF303099 GFP or DsRED strains. Nodule number was scored three and four weeks after inoculation. For phenotyping of *rpg-6*, WT and SL0181 (M4) plants were grown at 24 °C, 16 hours photoperiod as previously described (Groth et al., 2010). Nodules and uninfected nodule primordia were quantified 14 days after inoculation with DsRED-expressing *M. loti* MAFF303099 by fluorescence and bright-field microscopy (Leica MZ16 FA). The number of infection events was determined by microscopy of the whole root stained with 5-bromo-4-chloro-3-indolylbeta-D-galacto-pyranoside (X-Gal) at 4 and 10 dpi with *M. loti* R7A LacZ; at least nine plants were scored at each time point. LacZ staining, observation of GFP-marked *M. loti*-inoculated roots, and light microscopy of nodule sections were performed as previously described (Qiu et al., 2015).

For mycorrhizal analysis, *L. japonicus* seedlings were grown in pots containing sand and perlite (1:4) with sterile *R. irregularis* spores. The roots were stained with ink/vinegar and fungal structures quantified five weeks after inoculation as previously described (Zhang et al., 2015). The samples were analyzed at 10x magnification with a bright-field microscope (Nikon Eclipse). Images of roots stained with WGA-Alexa Fluor 488 were taken with confocal microscopy (Olympus FV10-ASW).

### Complementation tests

Roots of WT, *rpg-1* or *rpg-2* mutants were transformed with p*UB:RPG* using *A. rhizogenes* AR1193-mediated hairy root transformation. The transformed chimeric plants were transplanted into vermiculite/perlite pots and inoculated with *M. loti* R7A LacZ after five to seven days. Infection events were analyzed at seven dpi and the nodulation phenotypes were scored two or three weeks after inoculation.

### Gene expression pattern analysis

*L. japonicus* WT (Gifu), *rpg-1, rpg-2, nin-2*, and *ern1-2* seedlings were grown on FP agar medium for seven days. Plants were then either inoculated with *M. loti* R7A or 10 nM purified *M. loti* NFs was added. Samples were collected at 0, 1, 3, and 7 days after *M. loti* inoculation or 0, 6, 12, and 24 h after NF treatment, immediately frozen in liquid nitrogen, and stored at -80 °C until use. Total RNA was extracted using the TRIpure Isolation Reagent (Aidlab, China); RNA was reverse transcribed using TransScript one-step gDNA removal and cDNA synthesis SuperMix (Trans Gen Biotech). qRT-PCR reactions were performed with the TOYOBO SYBR Green Realtime PCR Master Mix (TOYOBO) and analyzed with a step-one Plus PCR system (ABI). *Lotus* Ubiquitin (Lj5g3v2060710.1) was used as a reference gene to normalize expression. All of the primers used for qRT-PCR of target transcripts are shown in Supplemental Table S1.

For promoter GUS assays, the p*RPG:GUS*, p*RPGΔS1:GUS*, p*RPGΔS2:GUS* construct was transferred into AR1193, then expressed in *L. japonicus* WT by hairy root transformation. Transgenic plants were transferred into a 1:1 vermiculite:perlite mixture and inoculated with *M. loti* R7A LacZ after five to seven days. GUS expression was analyzed at seven and 14 dpi.

### Electrophoresis mobility shift assays (EMSA)

NIN carrying a C-terminal His tag was purified as previously described (Xie et al., 2012). The RPG promoter regions S1(−1037 to −1231 bp), S2 (−141 to −339 bp), ΔS1, and ΔS2 were amplified via PCR using p*RPG:GUS*, p*RPGΔS1:GUS*, and p*RPGΔS2:GUS* as the template; primers are shown in Supplemental Table S1. PCR products were fluorescently labeled at the 5′ ends with Cy5 (Yingjun Corp. China) and purified by gel extraction (OMEGA Bio-TEK). Fluorescently-labeled DNA was then detected using a Biophotometer Plus (Eppendorf) and 1 nM of DNA incubated with the purified NIN protein in 20 μL of binding buffer (20 mm Tris, pH 7.5; 5% [w/v] glycerol; 10 mm MgCl2; 0.25 mm dithiothreitol; 0.8 μg bovine serum albumin [BSA]; and 1 μg salmon sperm DNA). After incubation at 30 °C for 20 min, the products were electrophoresed at 4 °C on a 6% native polyacrylamide gel in Tris-borate/EDTA buffer for 2 h at 100 V. Fluorescence in the gel was detected with a Starion FLA-9000 (FujiFilm).

### Dual-luciferase reporter assays in *N. benthamiana*

The dual-luciferase reporter assay was performed in *N. benthamiana* leaves as previously described (Luo et al., 2021). The indicated constructs were transferred into *A. tumefaciens* GV3101 (pSoup), then introduced into *N. benthamiana* leaves by infiltration. After two days, the LUC/REN ratio was measured with the dual-luciferase reporter assay system following the manufacturer’s protocols (Promega). Mean values and standard deviations were calculated from three biological replicates.

### Protein subcellular localization and co-localization in *N. benthamiana* leaves or *L. japonicus* root protoplasts

CERBERUS-mCherry and the other organelle markers used for protein subcellular localization analysis in *N. benthamiana* leaves have been described previously (Liu et al., 2021). The constructs were introduced into *A. tumefaciens* EHA105 by electroporation, and *N. benthamiana* leaves were infiltrated with the resulting strains either alone or together. All were infiltrated with p19, which inhibits gene silencing (Voinnet et al., 2003). Images were taken two days later with laser scanning confocal microscopy (Leica TCS SP8). The level of colocalization was analyzed using ImageJ. All protein subcellular localization assays were repeated at least three times.

For transient expression in *L. japonicus* root protoplasts, constructs (pA7-GFP-RPG, pA7-GFP-RPG, and ASTRAY*-*mRFP) were transiently expressed or co-expressed in *L. japonicus* root protoplasts using a DNA-PEG-calcium transfection method (Jia et al., 2018). Images were taken 16 h after transfection by laser scanning confocal microscopy (Leica TCS SP8). For GFP, the filter sets for excitation and emission were 488 nm and 498-550 nm, respectively; for mCherry, DsRed, and mRFP, they were 561 nm and 575-650 nm. The level of colocalization was analyzed using ImageJ. All protein subcellular localization assays were repeated at least three times.

### Whole-mount immunolocalization assays for RPG subcellular localization in *L. japonicus* roots

The p*35S:GFP-RPG* or p*RPG:GFP-RPG* plasmid was introduced into *A. tumefaciens* strain EHA105, then expressed in *rpg-1* by *A. tumefaciens-*mediated transformation (Tirichine et al., 2005) to generate stably transformed plants.

RPG subcellular localization was analyzed using whole-mount immunolocalization as previously described (Sauer et al., 2006). Briefly, transgenic plants were inoculated with MAFF303099/RFP and 5-7 days after inoculation, the roots were submerged in fixative solution (4% formaldehyde in phosphate-buffered saline (PBS)) in a vacuum desiccator for 1 h. Fixative solution was removed and seedlings were washed two times for 5-10 min each with 1x PBS at room temperature. This was followed by two washes with water for 5 min each. Root pieces were transferred to microscope slides and dried overnight, then root tissue was rehydrated by pipetting 1**×** PBS onto the microscope slides and incubating for 5 min at room temperature. Roots were collected into 2 mL EP tubes, then permeated with 2% Driselase in PBS and incubated for 60 min at 37 °C, followed by five washes with 1**×** PBS for 10 min each. A mixture of 3% IGEPAL CA-630 with 10% DMSO in PBS was added, then after 1 h the tissues were washed with 1**×** PBS five times for 10 min each. After blocking with 3% BSA in PBS, the fixed roots were incubated with primary antibody (anti-GFP, 1:300, Abmart) for at least 4 h at 37 °C. Alexa Fluor 488-conjugated AffiniPure Donkey Anti-Mouse IgG secondary antibody (1:500, Jackson) was added and incubated for at least 3 h at 37 °C, then samples were washed with 1**×** PBS five times for 10 min each. Images were taken with a confocal microscope (Leica TCS SP8). For RFP, the filter sets for excitation and emission were 561 nm and 575-650 nm, respectively; for Alexa 488, they were 488 nm and 498-519 nm.

### Protein-protein interaction assays

Interactions between CERBERUS, WD40, and ARM were assayed using the yeast two-hybrid system as previously described (Liu et al., 2021). The yeast strain AH109 was transformed with the constructs in destination vectors using lithium acetate transformation (Yeast Protocols Handbook PT3024-1, Clontech). The transformants were grown on synthetic defined medium (0.67% yeast nitrogen base, 2% Bacto-agar and amino acid mix) without the appropriate auxotrophic markers after gradient dilution. These assays were repeated three times.

For split-luciferase complementation imaging assays in *N. benthamiana* leaves, LUCc-RPG was co-expressed with CERBERUS-LUCn, ARM-LUCn, in *N. benthamiana* leaves *via* agroinfiltration with p19, which inhibits gene silencing. The transformed plants were grown in a growth chamber. After two days, images were captured by CCD (TANON 5200, China) after 1 mM luciferin (Promega) was sprayed onto the leaves. All images were acquired using the same exposure settings. Each interaction group was validated with at least three replicates, and two or three independent experiments were performed.

For co-immunoprecipitation (Co-IP) assays, GFP-RPG and CERBERUS-mCherry were co-expressed in *N. benthamiana* leaves. Leaves were harvested 60 h after agroinfiltration, and approximately 0.6 g of plant tissue was extracted with 2 ml lysis buffer (50 mM Tris-MES at pH 8.0, 0.5 M sucrose, 1 mM MgCl_2_, 10 mM EDTA, 5 mM DTT, 0.2% NP-40, 1 mM phenylmethanesulfonyl fluoride (PMSF] and proteinase inhibitor cocktail tablet (Roche]) for 15 min then centrifuged at 12,000 rpm for 10 min. The supernatants were collected for Co-IP. Samples were incubated for 1.5 h with 30 µL Anti-RFP Affinity beads 4FF (Cat. SA072C, SMART) at 4 °C on a rotating wheel, then centrifuged at 2000 rpm for 2 min at 4 °C. The beads were washed and analyzed by immunoblotting using anti-mCherry (Cat.T0090, Affinity Biosciences, Cincinnati, OH, USA) and anti-GFP antibody (Cat. M20004L, Abmart). Approximately 10 μL of lysis buffer containing total protein was loaded as the input control.

For BiFC assays, the construct p*AtUBI10:nVenus-CERBERUS/*p*LjUBI1:cVenus-RPG* was expressed in *N. benthamiana* leaves by agroinfiltration with p19. Transformed plants were grown in a growth chamber, and images were captured two to three days later by laser scanning confocal microscopy (Leica TCS SP8). The BiFC construct was also expressed in WT *L. japonicus* or *M. truncatula sunn-1* by hairy root transformation. The transgenic hairy roots were scored based on the NLS-DsRed marker and inoculated with *M. loti* MAFF303099/RFP or Sm1021/mCherry (OD_600_: 0.001). Images were analyzed at five to seven dpi. The filter sets for excitation and emission were 514 nm and 524 to 545 nm, respectively, for Venus, and 561 nm and 600 to 630 nm for DsRed. All BiFC experiments were repeated twice, and at least five leaves or roots were analyzed each time.

### Statistical analysis

Statistical significance was analyzed by Student’s *t-test* (\**P* < 0.05, \*\**P* < 0.01, \*\*\**P* < 0.001, \*\*\*\**P* < 0.0001) and error bars indicate SD. Histograms were generated using GraphPad Prism 8.0 software.

### Accession Numbers

Sequence data from this article can be found in the GenBank data libraries under accession number ON756094 for LjRPG.

## SUPPLEMENTAL INFORMATION

This manuscript contains 15 Supporting Figures and 2 Supporting Tables.

**Supplemental Figure S1**. Nodule sections of nodules formed on *L. japonicus rpg* mutants.

**Supplemental Figure S2**. *RPG* complementation of *rpg-1* and *rpg-2* mutants.

**Supplemental Figure S3**. Map-based cloning of the *rpg-6*.

**Supplemental Figure S4**. Infection and nodulation phenotypes of *rpg*::LORE1 insertion mutants.

**Supplemental Figure S5**. Nodule phenotype and complementation of *rpg-6*.

**Supplemental Figure S6**. *RPG* expression in *LORE1* insertion mutant roots after inoculation with *M. loti* R7A.

**Supplemental Figure S7**. Expression of early nodulin genes in wild type (WT) and *rpg* mutant roots after inoculation with *M. loti* R7A.

**Supplemental Figure S8**. Root colonization of *rpg* mutants by the AM fungus *R. irregularis*.

**Supplemental Figure S9**. Alignment of putative NIN-binding sites in the *RPG* promoter region.

**Supplemental Figure S10**. GFP-RPG can rescue *rpg-1* phenotype.

**Supplemental Figure S11**. Nodulation phenotype of the *rpg-1* mutant stably transformed with p*RPG:GFP-RPG*.

**Supplemental Figure S12**. Subcellular localization of GFP-RPG in *L. japonicus* root hairs after inoculation with *M. loti*.

**Supplemental Figure S13**. West blot analysis of full-length RPG and domains of RPG fused to GFP extracted from transiently transformed *N. benthamiana* leaves.

**Supplemental Figure S14**. RPG CC can rescue *rpg-1* nodulation phenotype.

**Supplemental Figure S15**. Co-localization of RPG with ER and TGN/EE markers but not Golgi or MVB markers.

**Supplemental Table S1**. Primers used in this study.

**Supplemental Table S2**. Constructs used in this study.

## ACKNOWLEDGMENTS

We thank Prof. Chi-Kuang Wen (CEMPS, CAS) and Prof. Yan Zhang (Shandong Agriculture U. China) for providing markers. Dr. Wenjuan Cai and Dr.Shuining Yin (CEMPS, CAS, China) for help with microscopy on this study. This work was funded by grants from CAS project for Young Scientists in Basic Research (YSBR-011), Program of Shanghai Academic/Technology Research Leader (21XD1403900), the Strategic Priority Research Program of the Chinese Academy of Sciences (XDB27040208), and the National Natural Science Foundation of China (31400214).

## AUTHOR CONTRIBUTIONS

X.L. Li and F. Xie designed the experiments, X.L. Li performed most of the experiments, M.X. Liu mapped *rpg-2*. M. Cai performed some promoter analysis. DC, MG and MP identified *rpg-6*. AH, MP, TLW and JAD contributed with *rpg-1, rpg-2* and *rpg-6* materials. X.L. Li, M.X. Liu and F. Xie analyzed the data and wrote the manuscript with editing by JAD.

## REFERENCES

Arrighi JF, Godfroy O, de Billy F, Saurat O, Jauneau A, Gough C (2008) The RPG gene of Medicago truncatula controls Rhizobium-directed polar growth during infection. Proc Natl Acad Sci USA 105: 9817–9822

Bar-Peled M, Raikhel NV (1997) Characterization of AtSEC12 and AtSAR1. Proteins likely involved in endoplasmic reticulum and Golgi transport. Plant Physiol 114: 315–324

Cerri MR, Wang Q, Stolz P, Folgmann J, Frances L, Katzer K, Li X, Heckmann AB, Wang TL, Downie JA, Klingl A, de Carvalho-Niebel F, Xie F, Parniske M (2017) The ERN1 transcription factor gene is a target of the CCaMK/CYCLOPS complex and controls rhizobial infection in Lotus japonicus. New Phytol 215: 323–337

DeBlasio SL, Luesse DL, Hangarter RP (2005) A plant-specific protein essential for blue-light-induced chloroplast movements. Plant Physiol 139: 101–114

Drakakaki G, van de Ven W, Pan S, Miao Y, Wang J, Keinath NF, Weatherly B, Jiang L, Schumacher K, Hicks G, Raikhel N (2012) Isolation and proteomic analysis of the SYP61 compartment reveal its role in exocytic trafficking in Arabidopsis. Cell Res 22: 413–424

Fahraeus G (1957) The infection of clover root hairs by nodule bacteria studied by a simple glass slide technique. J Gen Microbiol 16: 374–381

Fournier J, Timmers AC, Sieberer BJ, Jauneau A, Chabaud M, Barker DG (2008) Mechanism of infection thread elongation in root hairs of Medicago truncatula and dynamic interplay with associated rhizobial colonization. Plant Physiol 148: 1985–1995

Fukai E, Soyano T, Umehara Y, Nakayama S, Hirakawa H, Tabata S, Sato S, Hayashi M (2012) Establishment of a Lotus japonicus gene tagging population using the exon-targeting endogenous retrotransposon LORE1. Plant J 69: 720–730

Gage DJ (2004) Infection and invasion of roots by symbiotic, nitrogen-fixing rhizobia during nodulation of temperate legumes. Microbiol Mol Biol Rev 68: 280–300

Griesmann M, Chang Y, Liu X, Song Y, Haberer G, Crook MB, Billault-Penneteau B, Lauressergues D, Keller J, Imanishi L, Roswanjaya YP, Kohlen W, Pujic P, Battenberg K, Alloisio N, Liang Y, Hilhorst H, Salgado MG, Hocher V, Gherbi H, Svistoonoff S, Doyle JJ, He S, Xu Y, Xu S, Qu J, Gao Q, Fang X, Fu Y, Normand P, Berry AM, Wall LG, Ane JM, Pawlowski K, Xu X, Yang H, Spannagl M, Mayer KFX, Wong GK, Parniske M, Delaux PM, Cheng S (2018) Phylogenomics reveals multiple losses of nitrogen-fixing root nodule symbiosis. Science 361:125–126

Groth M, Takeda N, Perry J, Uchida H, Dräxl S, Brachmann A, Sato S, Tabata S, Kawaguchi M, Wang TL, Parniske M (2010) NENA, a Lotus japonicus homolog of Sec13, is required for rhizodermal infection by arbuscular mycorrhiza fungi and rhizobia but dispensable for cortical endosymbiotic development. Plant Cell 22: 2509–2526

Hirsch S, Oldroyd GE (2009) GRAS-domain transcription factors that regulate plant development. Plant Signal Behav 4: 698–700

Hossain MS, Liao J, James EK, Sato S, Tabata S, Jurkiewicz A, Madsen LH, Stougaard J, Ross L, Szczyglowski K (2012) Lotus japonicus ARPC1 is required for rhizobial infection. Plant Physiol 160: 917–928

Jia N, Zhu Y, Xie F (2018) An Efficient Protocol for Model Legume Root Protoplast Isolation and Transformation. Front Plant Sci 9: 670

Ke D, Fang Q, Chen C, Zhu H, Chen T, Chang X, Yuan S, Kang H, Ma L, Hong Z, Zhang Z (2012) The small GTPase ROP6 interacts with NFR5 and is involved in nodule formation in Lotus japonicus. Plant Physiol 159: 131–143

Kiss E, Oláh B, Kaló P, Morales M, Heckmann AB, Borbola A, Lózsa A, Kontár K, Middleton P, Downie JA, Oldroyd GE, Endre G (2009) LIN, a novel type of U-box/WD40 protein, controls early infection by rhizobia in legumes. Plant Physiol 151: 1239–1249

Kovács S, Kiss E, Jenei S, Fehér-Juhász E, Kereszt A, Endre G (2022) The Medicago truncatula IEF Gene Is Crucial for the Progression of Bacterial Infection During Symbiosis. Mol Plant Microbe Interact 35: 401–415

Lace B, Su C, Perez DI, Rodriguez-Franco M, Vernié T, Batzenschlager M, Egli S, Liu C-W, Ott T (2022) RPG acts as a central determinant for infectosome formation and cellular polarization during intracellular rhizobial infections. bioRxiv: 2022.2006.2003.494689

Li X, Zheng Z, Kong X, Xu J, Qiu L, Sun J, Reid D, Jin H, Andersen SU, Oldroyd GED, Stougaard J, Downie JA, Xie F (2019) Atypical Receptor Kinase RINRK1 Required for Rhizobial Infection But Not Nodule Development in Lotus japonicus. Plant Physiol 181: 804–816

Liu CW, Breakspear A, Guan D, Cerri MR, Jackson K, Jiang S, Robson F, Radhakrishnan GV, Roy S, Bone C, Stacey N, Rogers C, Trick M, Niebel A, Oldroyd GED, de Carvalho-Niebel F, Murray JD (2019a) NIN Acts as a Network Hub Controlling a Growth Module Required for Rhizobial Infection. Plant Physiol 179: 1704–1722

Liu CW, Breakspear A, Stacey N, Findlay K, Nakashima J, Ramakrishnan K, Liu M, Xie F, Endre G, de Carvalho-Niebel F, Oldroyd GED, Udvardi MK, Fournier J, Murray JD (2019b) A protein complex required for polar growth of rhizobial infection threads. Nat Commun 10: 2848

Liu J, Liu MX, Qiu LP, Xie F (2020) SPIKE1 Activates the GTPase ROP6 to Guide the Polarized Growth of Infection Threads in Lotus japonicus. Plant Cell 32: 3774–3791

Liu M, Jia N, Li X, Liu R, Xie Q, Murray JD, Downie JA, Xie F (2021) CERBERUS is critical for stabilization of VAPYRIN during rhizobial infection in Lotus japonicus. New Phytol 229: 1684–1700

Lombardo F, Heckmann AB, Miwa H, Perry JA, Yano K, Hayashi M, Parniske M, Wang TL, Downie JA (2006) Identification of symbiotically defective mutants of Lotus japonicus affected in infection thread growth. Mol Plant Microbe Interact 19: 1444–1450

Luo Z, Lin JS, Zhu Y, Fu M, Li X, Xie F (2021) NLP1 reciprocally regulates nitrate inhibition of nodulation through SUNN-CRA2 signaling in Medicago truncatula. Plant Commun 2: 100183

Maekawa T, Kusakabe M, Shimoda Y, Sato S, Tabata S, Murooka Y, Hayashi M (2008) Polyubiquitin promoter-based binary vectors for overexpression and gene silencing in Lotus japonicus. Mol Plant Microbe Interact 21: 375–382

Miyahara A, Richens J, Starker C, Morieri G, Smith L, Long S, Downie JA, Oldroyd GE (2010) Conservation in function of a SCAR/WAVE component during infection thread and root hair growth in Medicago truncatula. Mol Plant Microbe Interact 23: 1553–1562

Montiel J, Reid D, Grønbæk TH, Benfeldt CM, James EK, Ott T, Ditengou FA, Nadzieja M, Kelly S, Stougaard J (2021) Distinct signaling routes mediate intercellular and intracellular rhizobial infection in Lotus japonicus. Plant Physiol 185: 1131–1147

Murray JD, Karas BJ, Sato S, Tabata S, Amyot L, Szczyglowski K (2007) A cytokinin perception mutant colonized by Rhizobium in the absence of nodule organogenesis. Science 315: 101–104

Murray JD, Muni RR, Torres-Jerez I, Tang Y, Allen S, Andriankaja M, Li G, Laxmi A, Cheng X, Wen J, Vaughan D, Schultze M, Sun J, Charpentier M, Oldroyd G, Tadege M, Ratet P, Mysore KS, Chen R, Udvardi MK (2011) Vapyrin, a gene essential for intracellular progression of arbuscular mycorrhizal symbiosis, is also essential for infection by rhizobia in the nodule symbiosis of Medicago truncatula. Plant J 65: 244–252

Nelson BK, Cai X, Nebenführ A (2007) A multicolored set of in vivo organelle markers for co-localization studies in Arabidopsis and other plants. Plant J 51: 1126–1136

Newman-Griffis AH, Del Cerro P, Charpentier M, Meier I (2019) Medicago LINC Complexes Function in Nuclear Morphology, Nuclear Movement, and Root Nodule Symbiosis. Plant Physiol 179: 491–506

Nishimura R, Ohmori M, Kawaguchi M (2002) The novel symbiotic phenotype of enhanced-nodulating mutant of Lotus japonicus: astray mutant is an early nodulating mutant with wider nodulation zone. Plant Cell Physiol 43: 853–859

Oldroyd GE, Downie JA (2004) Calcium, kinases and nodulation signalling in legumes. Nat Rev Mol Cell Biol 5: 566–576

Oldroyd GE, Downie JA (2008) Coordinating nodule morphogenesis with rhizobial infection in legumes. Annu Rev Plant Biol 59: 519–546

Pajuelo E, Stougaard J (2005) Lotus japonicus’s a model system. In AJ Márquez, ed, Lotus japonicus Handbook. Springer Netherlands, Dordrecht, pp 3–24

Qiu L, Lin JS, Xu J, Sato S, Parniske M, Wang TL, Downie JA, Xie F (2015) SCARN a Novel Class of SCAR Protein That Is Required for Root-Hair Infection during Legume Nodulation. PLoS Genet 11: e1005623

Robertson JG, Warburton MP, Lyttleton P, Fordyce AM, Bullivant S (1978) Membranes in lupin root nodules. II. Preparation and properties of peribacteroid membranes and bacteroid envelope inner membranes from developing lupin nodules. J Cell Sci 30: 151–174

Rose A, Manikantan S, Schraegle SJ, Maloy MA, Stahlberg EA, Meier I (2004) Genome-wide identification of Arabidopsis coiled-coil proteins and establishment of the ARABI-COIL database. Plant Physiol 134: 927–939

Roy S, Liu W, Nandety RS, Crook A, Mysore KS, Pislariu CI, Frugoli J, Dickstein R, Udvardi MK (2020) Celebrating 20 Years of Genetic Discoveries in Legume Nodulation and Symbiotic Nitrogen Fixation. Plant Cell 32: 15–41

Saleh MC, van Rij RP, Hekele A, Gillis A, Foley E, O’Farrell PH, Andino R (2006) The endocytic pathway mediates cell entry of dsRNA to induce RNAi silencing. Nat Cell Biol 8: 793–802

Sauer M, Paciorek T, Benková E, Friml J (2006) Immunocytochemical techniques for whole-mount in situ protein localization in plants. Nat Protoc 1: 98–103

Schauser L, Roussis A, Stiller J, Stougaard J (1999) A plant regulator controlling development of symbiotic root nodules. Nature 402: 191–195

Schnabel E, Journet EP, de Carvalho-Niebel F, Duc G, Frugoli J (2005) The Medicago truncatula SUNN gene encodes a CLV1-like leucine-rich repeat receptor kinase that regulates nodule number and root length. Plant Mol Biol 58: 809–822

Schnabel E, Smith L, Long S, Frugoli J (2010) Transcript profiling in M. truncatula lss and sunn-1 mutants reveals different expression profiles despite disrupted SUNN gene function in both mutants. Plant Signal Behav 5: 1657–1659

Sinharoy S, Liu C, Breakspear A, Guan D, Shailes S, Nakashima J, Zhang S, Wen J, Torres-Jerez I, Oldroyd G, Murray JD, Udvardi MK (2016) A Medicago truncatula Cystathionine-β-Synthase-like Domain-Containing Protein Is Required for Rhizobial Infection and Symbiotic Nitrogen Fixation. Plant Physiol 170: 2204–2217

Soyano T, Hirakawa H, Sato S, Hayashi M, Kawaguchi M (2014) Nodule Inception creates a long-distance negative feedback loop involved in homeostatic regulation of nodule organ production. Proc Natl Acad Sci USA 111: 14607–14612

Suetsugu N, Higa T, Kong SG, Wada M (2015) PLASTID MOVEMENT IMPAIRED1 and PLASTID MOVEMENT IMPAIRED1-RELATED1 Mediate Photorelocation Movements of Both Chloroplasts and Nuclei. Plant Physiol 169: 1155–1167

Tirichine L, Herrera-Cervera JA, Stougaard J (2005) Transformation-regeneration procedure for Lotus japonicus. In AJ Márquez, ed, Lotus japonicus Handbook. Springer Netherlands, Dordrecht, pp 279–284

Truebestein L, Leonard TA (2016) Coiled-coils: The long and short of it. Bioessays 38: 903–916

Ueda T, Yamaguchi M, Uchimiya H, Nakano A (2001) Ara6, a plant-unique novel type Rab GTPase, functions in the endocytic pathway of Arabidopsis thaliana. Embo j 20: 4730–4741

Urbanski DF, Malolepszy A, Stougaard J, Andersen SU (2012) Genome-wide LORE1 retrotransposon mutagenesis and high-throughput insertion detection in Lotus japonicus. Plant J 69: 731–741

van Velzen R, Holmer R, Bu F, Rutten L, van Zeijl A, Liu W, Santuari L, Cao Q, Sharma T, Shen D, Roswanjaya Y, Wardhani TAK, Kalhor MS, Jansen J, van den Hoogen J, Gungor B, Hartog M, Hontelez J, Verver J, Yang WC, Schijlen E, Repin R, Schilthuizen M, Schranz ME, Heidstra R, Miyata K, Fedorova E, Kohlen W, Bisseling T, Smit S, Geurts R (2018) Comparative genomics of the nonlegume Parasponia reveals insights into evolution of nitrogen-fixing rhizobium symbioses. Proc Natl Acad Sci USA 115: E4700–E4709

Voinnet O, Rivas S, Mestre P, Baulcombe D (2003) An enhanced transient expression system in plants based on suppression of gene silencing by the p19 protein of tomato bushy stunt virus. Plant J 33: 949–956

Wang DY, Fulthorpe R, Liss SN, Edwards EA (2004) Identification of estrogen-responsive genes by complementary deoxyribonucleic acid microarray and characterization of a novel early estrogen-induced gene: EEIG1. Mol Endocrinol 18: 402–411

Wang JG, Li S, Zhao XY, Zhou LZ, Huang GQ, Feng C, Zhang Y (2013) HAPLESS13, the Arabidopsis μ1 adaptin, is essential for protein sorting at the trans-Golgi network/early endosome. Plant Physiol 162: 1897–1910

Weber E, Engler C, Gruetzner R, Werner S, Marillonnet S (2011) A modular cloning system for standardized assembly of multigene constructs. PLoS One 6: e16765

Xie F, Murray JD, Kim J, Heckmann AB, Edwards A, Oldroyd GE, Downie JA (2012) Legume pectate lyase required for root infection by rhizobia. Proc Natl Acad Sci USA 109: 633–638

Yano K, Shibata S, Chen WL, Sato S, Kaneko T, Jurkiewicz A, Sandal N, Banba M, Imaizumi-Anraku H, Kojima T, Ohtomo R, Szczyglowski K, Stougaard J, Tabata S, Hayashi M, Kouchi H, Umehara Y (2009) CERBERUS, a novel U-box protein containing WD-40 repeats, is required for formation of the infection thread and nodule development in the legume-Rhizobium symbiosis. Plant J 60: 168–180

Yokota K, Fukai E, Madsen LH, Jurkiewicz A, Rueda P, Radutoiu S, Held M, Hossain MS, Szczyglowski K, Morieri G, Oldroyd GE, Downie JA, Nielsen MW, Rusek AM, Sato S, Tabata S, James EK, Oyaizu H, Sandal N, Stougaard J (2009) Rearrangement of actin cytoskeleton mediates invasion of Lotus japonicus roots by Mesorhizobium loti. Plant Cell 21: 267–284

Zhang D, Aravind L (2010) Identification of novel families and classification of the C2 domain superfamily elucidate the origin and evolution of membrane targeting activities in eukaryotes. Gene 469: 18–30

Zhang X, Pumplin N, Ivanov S, Harrison MJ (2015) EXO70I Is Required for Development of a Sub-domain of the Periarbuscular Membrane during Arbuscular Mycorrhizal Symbiosis. Curr Biol 25: 2189–2195

